# A Multimodal Atlas Reveals the Anatomical Distribution of Medium Spiny Neuron Subtypes and a Novel RGS6+ Population in the Primate Striatum

**DOI:** 10.1101/2025.11.16.688724

**Authors:** Ghada Abdelhady, Olivia R. Brull, Jing He, Meng K. Lin, Adriana Galvan, Andreas R. Pfenning, Andreea C. Bostan, William R. Stauffer

## Abstract

The primate striatum and its principal neuron type, the medium spiny neuron (MSN), integrate cortical and subcortical signals related to movement, cognition, and emotion. These signals are processed through cell type specific circuits traditionally defined by MSN dopamine receptor expression. However, classification by dopamine receptor type alone fails to fully specify MSN diversity and falls short of capturing the functional complexity of the striatum. Here, we combined single-nucleus multi-omic sequencing and high-plex spatial transcriptomics to build a comprehensive atlas of MSNs in the macaque striatum. Using multi-omic sequencing, we profiled MSNs across four anatomically and functionally defined territories, and we mapped these subtypes back into their anatomical context by integrating the multi-omic data with ∼5.4 million spatially resolved cells sampled across the full rostral-caudal and dorsal-ventral extent of the striatum. This approach revealed two previously undocumented ventral striatum (VS) subtypes, D1-VS-RGS6 and D2-VS-RGS6, which are molecularly distinct from known ventral striatal MSNs yet share core limbic features. We also uncovered gradients in matrix-compartment cell types along the rostral-caudal axis. Finally, by integrating MSN subtype-specific transcriptomes and ATAC-seq-derived regulatory annotations with human GWAS data, we demonstrate strong, cell-type-specific enrichment of polygenic risk for Parkinson’s disease, substance use disorders, and psychiatric and cognitive traits, including a striking association of D2-VS-RGS6 with schizophrenia and bipolar disorder. Together, this multimodal atlas provides a foundation for linking primate striatal cell types to circuit function and disease mechanisms.

**HIGHTLIGHTS:**

1. Multimodal analysis of NHP striatum reveals heterogeneous cell type distribution
2. Two previously uncharacterized MSN subtypes in the ventral striatum express RGS6
3. Ventral striatum cell types exhibit similar characteristics across the Rostro-Caudal axis
4. NHP cell types show strong, cell type specific associations to genomic disease predictors

## INTRODUCTION

The basal ganglia are crucial for learning,^1^ movement,^2,3^ and cognition,^4–6^ and their dysfunction is implicated in movement disorders such as Parkinson’s disease (PD),^7^ psychiatric diseases including obsessive compulsive disorder,^8^ and nearly all substance use disorders.^9^ This broad functional diversity arises from cell type specific circuitry centered on the striatum - the main basal ganglia input nucleus - and its predominant neuronal population, Medium Spiny Neurons (MSNs). Among the most studied of these cell type specific circuits are the Cortico–Basal Ganglia–Thalamic–Cortical (CBGTC) loops, which connect the basal ganglia with nearly all cortical regions.^10^ Corticofugal fibers forming these circuits project to the striatum and synapse onto MSNs.^11^ From there, signals are split into two pathways characterized by two MSN types: MSNs expressing type 1 dopamine receptors (D1-MSNs) that project to the internal segment of the globus pallidus (GPi) or the *substantia nigra pars reticulata* (SNpr) and form the “direct” pathway, and MSNs expressing type 2 dopamine receptors (D2-MSNs) that project to the external segment (GPe) and form the “indirect” pathway.^12–14^ Although the majority of research focuses on the direct and indirect pathways, the canonical MSN types that give rise to them represent only a portion of the cellular and circuit heterogeneity within the striatum. For example, D1-expressing MSNs located in the striosome compartment do not participate in the direct pathway but instead project to midbrain dopamine neurons via distinct circuits that likely subserve different behavioral functions.^15^ Thus, classification by dopamine receptor type alone underspecifies MSN diversity, and a more complete understanding of the phenotypic and anatomical diversity of MSNs—and their relationship to specific functional circuits—is essential for elucidating circuit-based mechanisms of behavior and disease.

Single-cell atlases that integrate genomic, epigenomic, and spatial modalities form the foundation for understanding brain function across multiple levels. Such datasets enable the dissection of cell type specific neural circuits,^16,17^ the identification of molecular determinants of human disease,^18–20^ and the generation of insights into evolution of brain systems.^21^ Despite the unmatched utility of nonhuman primates (NHP) for understanding human brain function,^22^ and the history of NHP research in the development of treatments for PD,^7,23–25^ a comprehensive multimodal atlas encompassing the large and multipotent NHP striatum remains unavailable. Transcriptomic data have provided initial estimates of the cell types in the NHP striatum,^26–28^ but these have mostly focused on the rostral striatum. Neurochemical markers such as calbindin (CB), substance P (SP), and acetylcholinesterase (AChE) have been particularly useful in defining the ‘neostriatal mosaic’ of striosome and matrix,^29–31^ but this has proven more challenging in the ventral striatum (VS), especially in the caudal portion that interfaces with the amygdala.^32^ An approach combining single cell genomics and spatial transcriptomics can provide new insights into this brain structure, but the large size and complicated geometry of the primate striatum present formidable challenges.

Here, we address these challenges with novel computational approaches that combine single cell multi-omics with a new large-scale spatial transcriptomics dataset. By sampling cells and tissue along both the rostral-caudal (RC) and dorsal-ventral (DV) axes of the macaque striatum, we identify previously unknown MSN subtypes. By using spatial transcriptomics and mapping these cell types throughout the striatum, we demonstrate that these cell types are not homogeneously distributed, but rather reflect the functional complexity of the striatum. This study, therefore, fills a critical gap by providing a comprehensive atlas of MSN subtypes in the NHP striatum that demonstrates how the anatomical distribution of MSN subtypes relates to the functional diversity of the striatum.

## RESULTS

### Integrated multi-omic sequencing and spatial transcriptomic analysis of the NHP striatum

To map the molecular properties of MSNs across the striatum, we integrated single-cell sequencing with spatial transcriptomics, applying these methods to distinct functional territories of the Old-World monkey striatum. For single cell sequencing, we harvested striatal tissue from rhesus macaque monkeys (Macaca mulatta) (n=5, 3 females and 2 males, age range at perfusion 3-12 years). We sampled four regions in the rostral-caudal (RC) and dorsal-ventral (DV) planes. These samples included the associative striatum, nucleus accumbens (NAc), sensorimotor putamen, and caudal ventral putamen (PUTv) (Figure 1A). We dissociated the samples to single nuclei and performed single nucleus RNA sequencing (snRNA-Seq)(n = 2) or multi-omic sequencing, including snRNA-Seq and single nucleus Assay for Transposase-Accessible Chromatin with sequencing (snATAC-Seq)(n=3). After quality control filtering, the dataset comprised **158,728 nuclei** that represented all major cell classes (Supplementary Figure 1C,D). We separated MSNs from glia and interneurons, which yielded **39,352 MSNs**. By stratifying MSNs into subtypes, we discovered eleven distinct MSN or MSN-like subtypes, nine of which we previously described, including the MSN-like D1-NUDAPS and D1-ICjs (Figure 1B).^26^ We noted substantial regional variations between MSN subtype composition. Low dimensionality projections of cells divided by region suggested that subtype similarity was determined by the RC axis to a greater degree compared to the DV axis (Figure 1C, top). This RC gradient was evident in both the dorsal and ventral territories; in particular, the PUTv exhibited a transcriptional profile more similar to that of the limbic NAc than to the sensorimotor putamen (Supplementary Figure 1A-B, Figure 1C, bottom). Although some of these regional differences are well known, some were surprising, and so we used spatial transcriptomics to develop a comprehensive understanding of the cell type structure of the NHP striatum.

**Figure 1.**
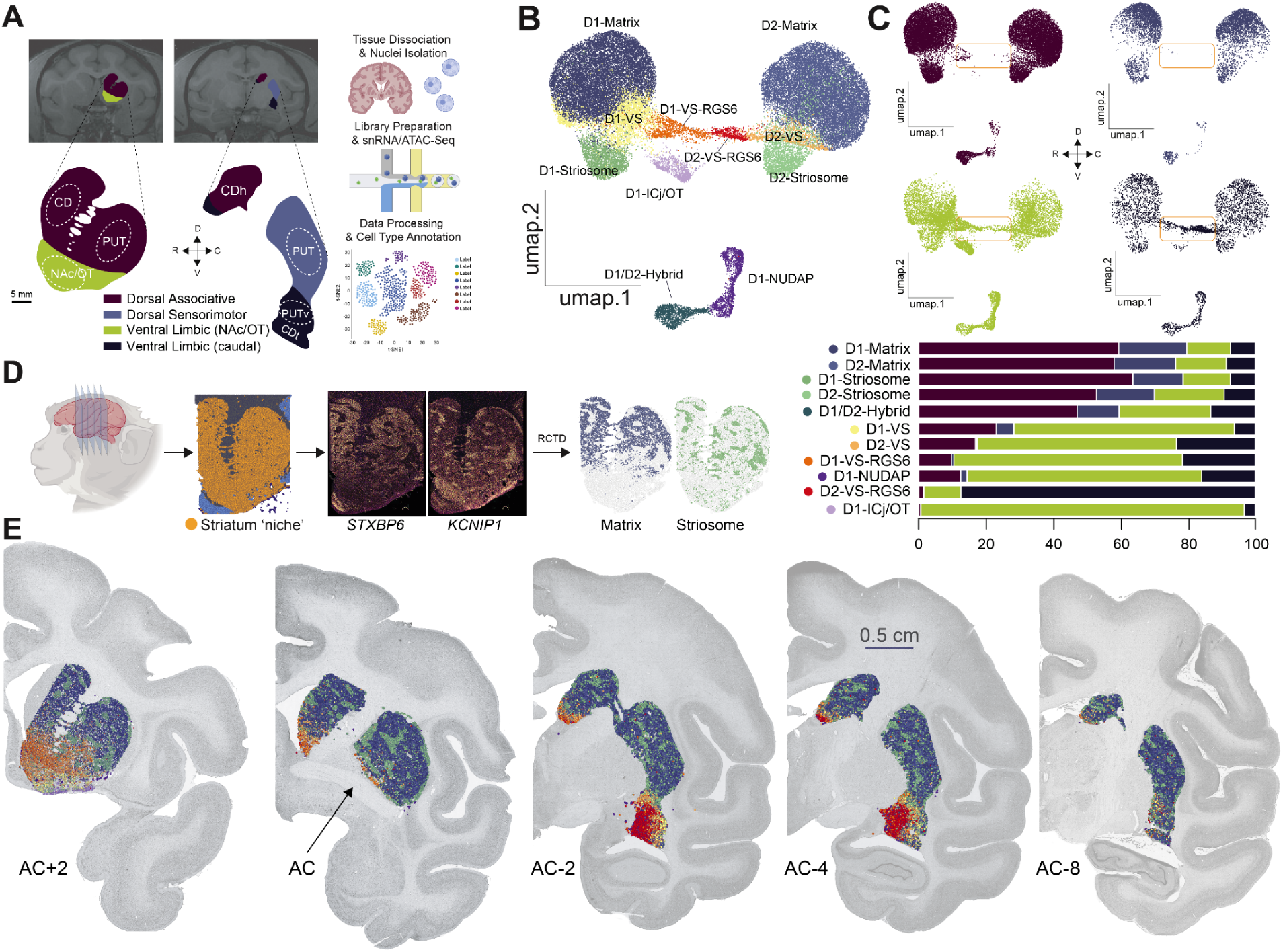
Mapping Cell Type Heterogeneity Across The Striatum. A. Sequencing workflow schematic diagram illustrating the tissue dissection at 2 key levels across the rostral-caudal axis in rhesus macaques (N=5), followed by snRNA-seq or snmulti-omic-seq of striatal tissue. Location of dissected tissue is shown in dotted outline. Some of the schematics are created in <u>BioRender.com</u>. B. Low dimensionality projection of MSNs (UMAP), after quality-control filtering, integration, re-clustering and cell subclasses annotation. Color assignments are consistent across all panels to aid visual comparison between figures. C. Top: Low dimensionality projection of MSNs split by the four striatal regions across R-C and D-V axes (UMAP), after QC filtering, integration of striatal MSNs nuclei from 4 striatal regions and 5 rhesus macaques and clustering. Four regions were annotated based on functional profiles into associative striatum, NAc, sensorimotor putamen, and PUTv. Orange box highlights MSN clusters overrepresented in ventral striatal regions. Bottom: Stacked barplot showing the relative regional proportions per cell type for the four functionally defined striatal regions, highlighting the predominance of D1-VS-RGS6 in the NAc and D2–VS-RGS6 in the PUTv. D. High-Plex Spatial Transcriptomics workflow schematic illustrating the tissue dissection at 5-7 levels across rostral-caudal axis in rhesus macaque (N=2) and in cynomolgus macaque (N=3), followed by spatial mapping of midbrain tissue, demarcation of striatal tissue using spatial niche assay, experimental checkpoint using mutually exclusive striatal compartments, STXBP6 for matrix and KCNIP1 for striosomes, RCTD cell type label prediction and computational checkpoint for predicted matrix and striosome cell types. E. Spatial cluster plot of the predicted cell types for five tissue sections across the R-C axis using the anterior Commissure (AC) as the reference level (AC+2, AC, AC-2, AC-4, AC-8). This highlights spatial patterns and gradients of cell types. Predictions of label annotations were implemented using our high quality integrated macaque striatum snRNA-Seq dataset as reference (in Figure 1B-C). Color assignments are consistent across all panels to aid visual comparison between figures.

For spatial transcriptomics, we used marker genes from snRNA-Seq to design a custom probe panel to detect known and novel MSN subtypes. In both rhesus and cynomolgus macaques (n=5, 2 females and 3 males, age range at perfusion 4-13 years), we cut coronal brain sections spanning ∼1 cm in the RC axis - from approximately 2 mm anterior to the anterior commissure (AC) to approximately 8 mm beyond the AC - and performed high-plex spatial transcriptomics. This enabled us to perform high-resolution, spatially-resolved cellular profiling (Figure 1D). After quality control filtering, we recovered **3,605,984** and **1,783,490** cells from rhesus and cynomolgus macaque, respectively, representing 99% of the total number of detected cells. We used anatomical landmarks to demarcate the boundaries of the striatum, which aligned well with the high spatial density of *DRD1* and *DRD2* transcripts in the striatum. We validated these boundaries with a ‘spatial niche assay’, which uses clusters or initial class annotations to identify cells with similar microenvironments based on the composition of spatially adjacent cell types (Supplementary Figure 3, Methods).

To align the MSN subtype annotations between snRNA-Seq and spatial transcriptomics data, we adapted Robust Cell Type Decomposition (RCTD)^33^, a computational approach for deconvolving spatial data using snRNA-seq reference data (Figure 1D). Here, RCTD-label transfer used the snRNA-Seq data set as a reference and determined, for each cell, the probabilities associated with each subtype label (Methods). We set a strict threshold, 0.5, and then assigned each cell the label with the probability that exceeded that threshold. If no label had a probability greater than 0.5, we designated the cell as unclassified. This process resulted in **849,315** confidently labeled cells, including **469,761 MSNs.** The MSN fraction constituted 56.5% of the total cell population and 92.6% of neurons. These numbers are consistent with prior estimations of striatal cell type densities and indicate that our samples are representative of NHP striatum.^34,35^ After annotation with RCTD-label transfer, we visualized cell types in their anatomical contexts, revealing clear separations between matrix, striosome, and VS compartments along the rostral-caudal (RC) axis of the NHP striatum (Figure 1E). Altogether, these results provide a comprehensive MSN map of the nonhuman primate striatum.

### Novel RGS6-Expressing MSN Subtypes are Characteristic of an Extended Limbic Striatum

We identified two novel clusters in the VS: *DRD1*- and *DRD2*-expressing MSNs that co-expressed *RGS6* (Regulator of G protein signaling 6), which we denote as D1-VS-RGS6 and D2-VS-RGS6, respectively (Figure 2A). To determine if these clusters represented distinct MSN subtypes, relative both to one another and to the nine previously described NHP MSN subtypes, we compared their gene expression, differential chromatin accessibility, and anatomical distributions. First, we identified marker genes for these two cell types (Figure 2B, Supplementary table S1). *RGS6* was a highly specific marker for both cell types. *DRD1* and *TAC3* were amongst the D1-VS-RGS6 markers, whereas *PENK* and *MOXD1* were abundant in D2-VS-RGS6. Next, we computed differential gene expression between each pair of cell types and used the count of significantly differentially expressed genes (DEGs) as a proxy for the degree of relatedness between cell types. The *RGS6* subtypes possessed hundreds of DEGs that set them apart from each other and the previously described VS MSNs.^26^ In fact, approximately 400 DEGs separated both D1-VS-RGS6 from D1-VS and D2-VS-RGS6 from D2-VS (Figure 2C). In comparison, D1-Matrix and D2-Matrix MSNs - widely acknowledged to be distinct cell types - were separated from one another by approximately 100 DEGs. We observed a similar pattern in chromatin accessibility, whereby differentially accessible chromatin regions reflected distinct MSN subtypes (Figure 2D). These multi-omic results clearly distinguish the *RGS6* subtypes from one another and the previously described MSN subtypes. These results indicate that D1-VS-RGS6 and D2-VS-RGS6 constitute two molecularly distinct MSN phenotypes.

**Figure 2.**
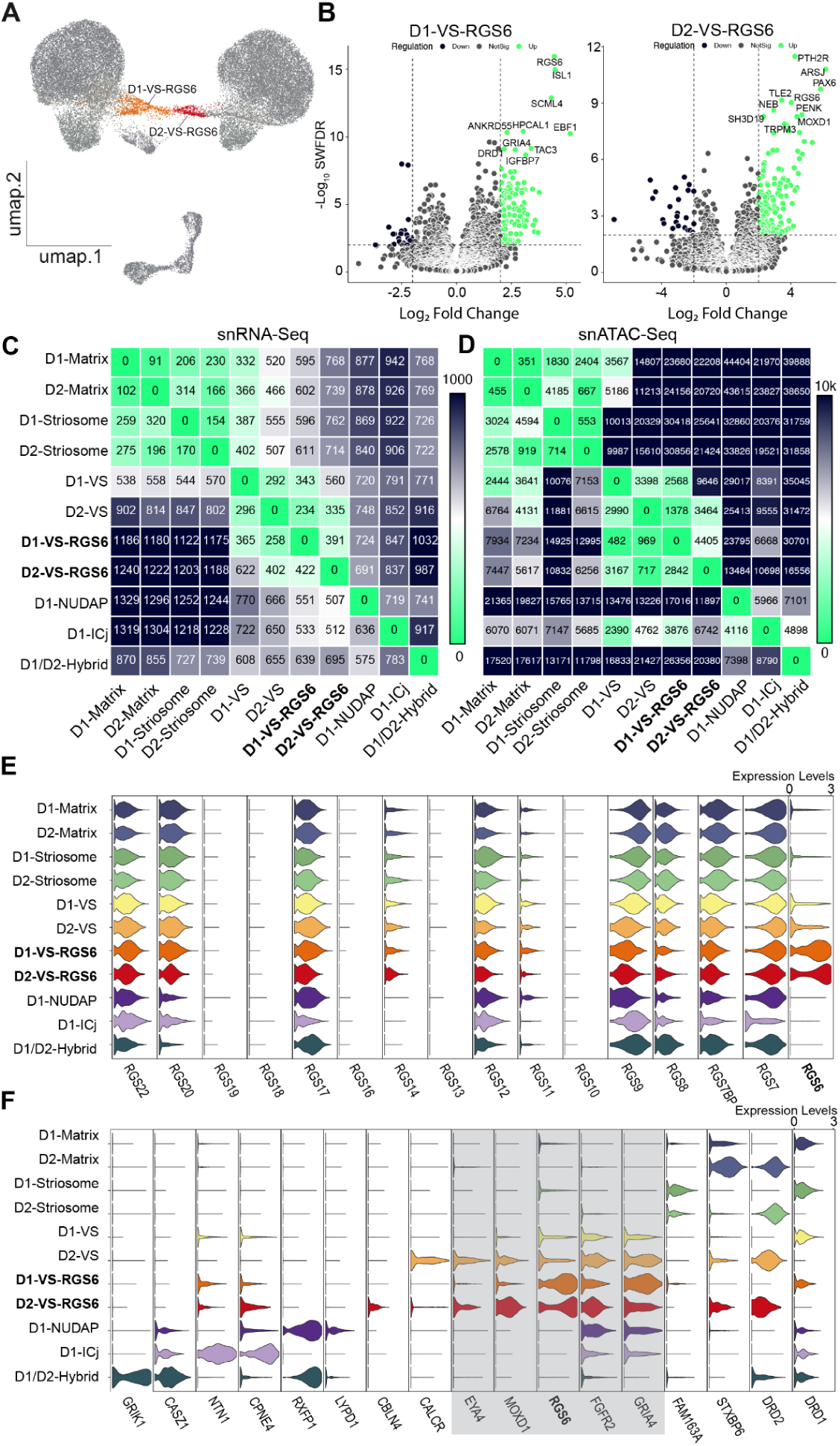
Identification of Novel RGS6-Expressing MSN Subtypes. A. Low dimensionality projection of MSNs (UMAP) highlighting D1-VS-RGS6 and D2-VS-RGS6. B. Differential gene expression analysis for D1-VS-RGS6 and D2-VS-RGS6 in contrast to all other MSN subtypes. x-axis: Log2(Fold Change) and y-axis: negative Log10(Weighted FDR). Top 10 genes for each cell type are annotated. C. Heatmap displaying the number of significantly upregulated differentially expressed genes (DEGs) between each pair of cell types, with lower number of DEGs indicating similarity. D. Heatmap displaying the number of significantly differential open chromatin regions (OCRs) between each pair of cell types, with lower number of differential OCRs indicating similarity. E. Violin plots for RGS gene family expression for the 11 cell subtypes of MSNs. Normalized gene expression per cell type is averaged across regions and animals. F.Violin plots for the MSNs cell type-specific marker genes used to identify and annotate the 11 MSNs subtypes. Shaded box highlighting ventral striatum markers: *GRIA4*, *FGFR2*, *RGS6*, *MOXD1* and *EYA4*. Normalized gene expression per cell type is averaged across regions and animals.

Regulators of G protein signaling (RGS) are GTPase-activating proteins that play a critical role in neurotransmission by regulating the duration and magnitude of G protein coupled receptor signaling.^36^ Thus, we wondered whether other RGS proteins have cell type specific functions in the striatum. Upon profiling the expression of 24 RGS-related genes across the 11 MSN subtypes, we noted non-specific expression of *RGS7*, *7BP*, *8*, *9*, *12*, *17*, *20* and *22*. Only *RGS6* exhibited a robust and cell type specific signal for any MSN subtype (Figure 2E). These results suggest that *RGS6* may play important and functionally specific roles in striatal signalling.

The differential expression and chromatin accessibility indicated that D1-VS-RGS6 and D2-VS-RGS6 were indeed distinct molecular phenotypes. Nevertheless, they shared many marker genes with the major limbic MSN subtypes D1-VS and D2-VS. These markers included *GRIA4* and a novel VS marker gene: fibroblast growth factor receptor 2 (*FGFR2*). *MOXD1* expression was found in all major VS MSNs, whereas *EYA4* emerged as a strong marker for both D2-VS and D2-VS-RGS6 subtypes (Figure 2F). This suggests that, although distinct, D1-VS-RGS6 and D2-VS-RGS6 share limbic characteristics with the other major VS MSN subtypes.

Spatial transcriptomic analysis of the NHP striatum revealed that D1-VS-RGS6 and D2-VS-RGS6, as well as the other two major VS MSN subtypes, were broadly distributed across the RC axis, but each subtype possessed unique spatial characteristics (Figure 3A-D). In the rostral striatum, the D1-VS-RGS6 population was enriched in the medial aspect of the NAc (Figure 3A), whereas the D1-VS population was restricted to the lateral shell region (Figure 3B, left). At this same RC level, the D2-VS-RGS6 population was almost nonexistent, but the D2-VS population was abundant and largely overlapping with the D1-VS-RGS6 population. There were few VS cell types located at the level of the AC, and they were all clustered in the medial caudate (CD) nucleus. Caudal to the AC, the D1-VS-RGS6 subtype was present in the medial caudate head (CDh), whereas the D2-VS-RGS6 subtype was highly enriched in the PUTv and CD tail (CDt). Thus, it appeared that D1-VS-RGS6 was present throughout the RC axis of the striatum, whereas D2-VS-RGS6 was only present in the caudal striatum. This was confirmed by correlation of the snRNA-Seq data with rostral and caudal spatial transcriptomic sections. For D1-VS-RGS6, the correlations between the snRNA-seq data and both rostral and caudal spatial transcriptomic sections were high, indicating the presence of the D1-VS-RGS6 subtypes in both rostral and caudal sections (Figure 3A, insets, R^2^ = 0.46 and 0.50, p = 5.5 x 10^-10^ and 6.7 x 10^11^, for rostral and caudal sections, respectively). In contrast, for D2-VS-RGS6, correlation between the snRNA-seq reference and rostral sections was low, but the correlation between the snRNA-seq reference and gene expression in the caudal striatum was high (Figure 3B, insets, R^2^ = 0.26 vs. 0.55, p = 7.8 x 10^-5^ vs 9.3 x 10^-12^, for rostral vs caudal sections). These results confirm that the D1-VS-RGS6 subtypes were present throughout the RC axis, but that the D2-VS-RGS6 subtypes were preferentially enriched in the caudal striatum. The D1-VS and D2-VS subtypes were also enriched in PUTv, but in more lateral territories, compared to D2-VS-RGS6. These results provide evidence for the limbic nature of the VS and reveal that it is characterized by the distinct spatial profiles of four major MSN subtypes.

**Figure 3.**
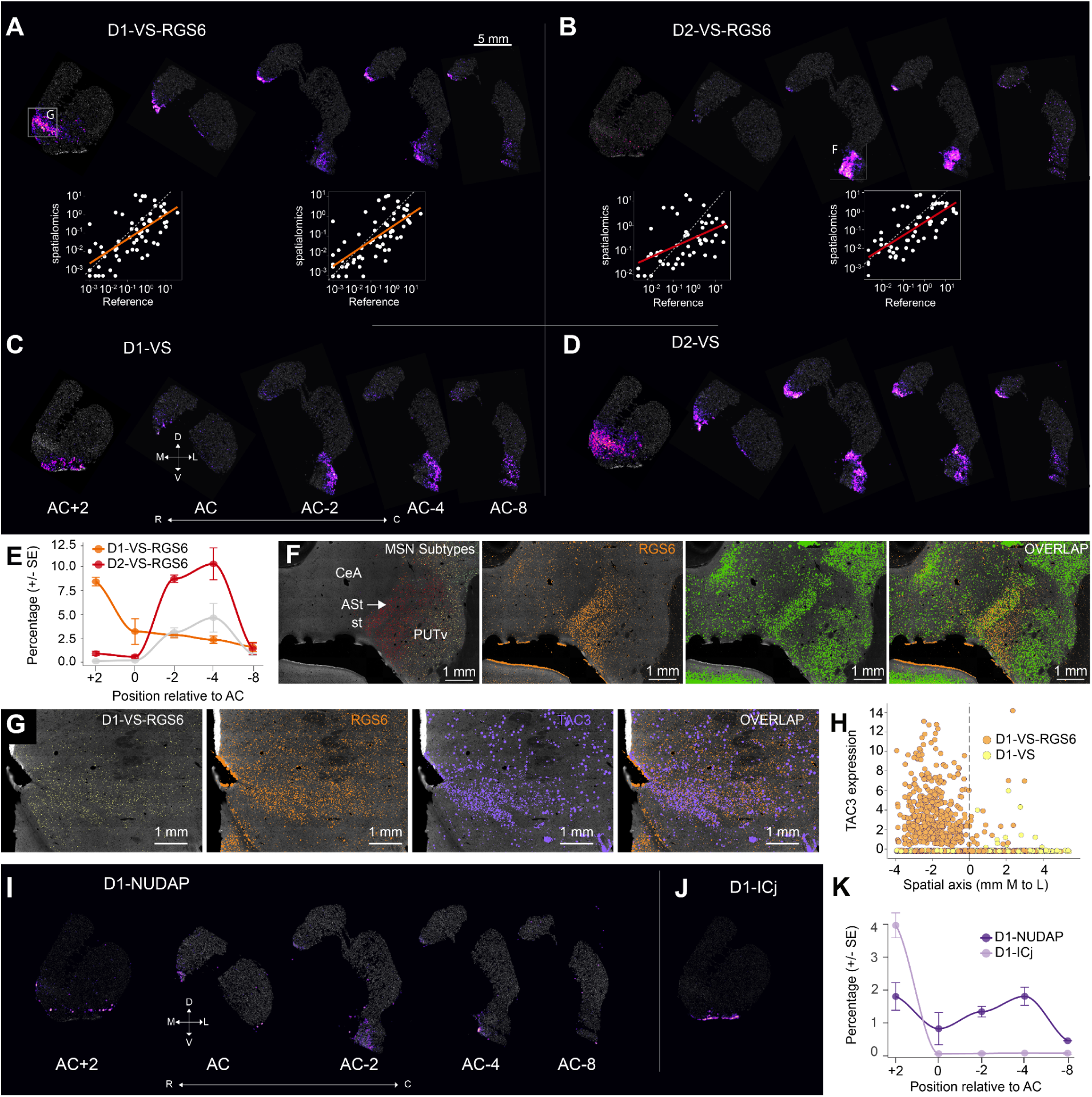
Spatial Patterning of Ventral Striatum Medium Spiny Neurons Across Rostral-Caudal Axis. A-D. Cell type density map highlighting differential spatial localization of D1-VS-RGS6, D2-VS-RGS6, D1-VS and D2-VS across 5 levels along the RC axis. A-B, Bottom: Evaluation of the mean gene expression correlation between the spatial query and the snRNA-Seq reference per cell type. Scatterplots are shown for rostral section (AC+2) and caudal section (AC-4) illustrating the increase in correlation in gene expression for D2-VS-RGS6 in the caudal section. E. Regional-cell type relative abundance gradient scatter plot smoothed using Locally Estimated Scatterplot Smoothing (LOESS) across the rostral-caudal axis. For each section, counts of D1-VS-RGS6 and D2-VS-RGS6 subtypes were aggregated across subjects and their relative proportions were calculated as to percentages. Fold change of D2-VS-RGS6 relative to D1-VS-RGS6 is plotted in grey. Error bars represent ± standard error of the mean (SEM) across sections within each region. F. Cell type density map for RGS6+ neurons (panel 1) and transcript density map for spatial localization of *RGS6*, *CALB1* and overlap (panel 2-4), highlighting RGS6+ calbindin poor region and RGS6+ calbindin rich region in the caudal ventral striatum. G. Cell type density map for D1-VS-RGS6 (panel 1) and transcript density map for spatial localization of *RGS6*, *TAC3* and overlap (panel 2-4) in the rostral ventral striatum. H. Regression discontinuity analysis highlighting the discrete transition in *TAC3* expression (y-axis) across a spatial boundary (x-axis=0) between D1-VS and D1-VS-RGS6. I. Cell type density map highlighting differential spatial localization of D1-NUDAP across 5 levels along the RC axis. K. Regional-cell type relative abundance gradient scatter plot smoothed using LOESS across the rostral-caudal axis. For each section, counts of D1-NUDAP and D1-ICj subtypes were aggregated across subjects and their relative proportions were calculated as percentages. Error bars represent ± SEM across sections within each region.

The appearance of the D2-VS-RGS6 subtype in the postcommissural striatum is striking. In fact, in these caudal sections, the D2-VS-RGS6 subtype had up to a 5-fold increase relative to D1-VS-RGS6 (Figure 3E). As described above, the majority of the D2-VS-RGS6 subtype was located in PUTv and CDt. Spatial niche analysis included the amygdalo-striatal transition zone as part of the striatum, indicating that this zone, from a genomic perspective, was similar to the striatum. We found that the D2-VS-RGS6 subtype was especially enriched there (Figure 3F, left). A closer look at *RGS6* and *CALB1* expression, the latter of which strongly marks the amygdalostriatal region, confirms this spatial relationship (Figure 3E). This result suggests that the D2-VS-RGS6 subtype could act as an interface between striatum and amygdala.

As noted in Figure 2B, the D1-VS-RGS6 cell type exhibited high expression of the neurokinin-B receptor *TAC3*. In our previous analysis in the rostral striatum, we identified a D1-VS archetype that was enriched in *TAC3*. ^26^ This convergence prompted a re-examination of the D1-VS and D1-VS-RGS6 clusters to determine whether *TAC3* was enriched in both subtypes. Our spatial transcriptomic data showed high correlation between the D1-VS-RGS6 subtype and expression of *TAC3* (Figure 3G). We used the medial-lateral axis in this section, which separates D1-VS-RGS6 from D1-VS, and demonstrated a discontinuity in *TAC3* expression between the laterally located D1-VS and medial D1-VS-RGS6 subtypes (Figure 3H, p = 6.4 x 10^-12^, regression discontinuity test). Thus, we conclude that the previously reported *TAC3* archetype is, in fact, the distinct D1-VS-RGS6 subtype.

We uncovered additional cell type specific and gene expression markers of the limbic striatum across the RC axis. First, we examined the distribution of D1-NUDAP cells, a prominent feature of the NAc.^26^ We discovered that D1-NUDAP islands were detectable throughout the RC axis (Figure 3I). In contrast, the D1-ICj islands, another prominent feature of the NAc, were only located in the rostral sections and always in close proximity to the olfactory tubercle (Figure 3J). These results, too, were highly reproducible across animals (Figure 3F). Altogether, the data indicate that PUTv is a caudal extension of the limbic striatum rather than a ventral extension of sensorimotor territories. Notably, the proportions of both D1- and D2-VS-RGS6 subtypes as well as D1-NUDAPs markedly decreased at the most caudal regions of the striatum (Figure 2A-C, Right). At this level, approximately 8-10 mm anterior to the intra-aural line, there is no detectable amygdala. We suspect that this marks the caudal boundary of the limbic striatum.

### Matrix compartment shows a D2-MSN bias in the Caudal Striatum

In the dorsal striatum, the spatial transcriptomic analysis revealed interesting cell type heterogeneity along the R-C axis (Figure 1F). Prior research has demonstrated that striosomes are a feature of the rostral striatum and decrease in the caudal direction.^31^ At the level of cell types, this trend was evident but minimal: we observed distinct striosomes even in the most caudal regions (Figure 4A). The caudal decrease in striosome cell types (Figure 4B) was mostly accounted for by an overabundance of D1- and D2-Striosome MSNs in the most rostral section, compared to all other sections. The matrix compartment was more interesting. There was a relative increase in the matrix cell types percentage across sections from the rostral to caudal direction. This increase was driven by cell type specific and spatial factors. The percentage of D1-Matrix MSNs was relatively constant along the RC axis. In contrast, the percentage of D2-Matrix MSNs increased significantly along the RC axis (Figure 4C, **D1-Matrix**: β=-0.38+0.24, p = 0.133 and **D2-Matrix**: β=1.44+0.3, p = 0.0005, linear mixed model (LMM) with region as a fixed effect and species as a random effect). Moreover, this relative increase appeared in the PUTv, beyond the posterior extent of the amygdala, where we suspect the limbic territory ends. This region was mostly devoid of D1-Matrix MSNs (Figure 4D, top right, oval), but contained an abundance of D2-Matrix MSNs (Figure 4D, bottom right, oval). This cell type specific spatial discrepancy was present in every animal we tested (Figures 4E,F). These results demonstrate that there is a gradient in cell type specific percentages that is caused mostly by an increase in D2-Matrix MSNs in the most caudal PUTv.

**Figure 4.**
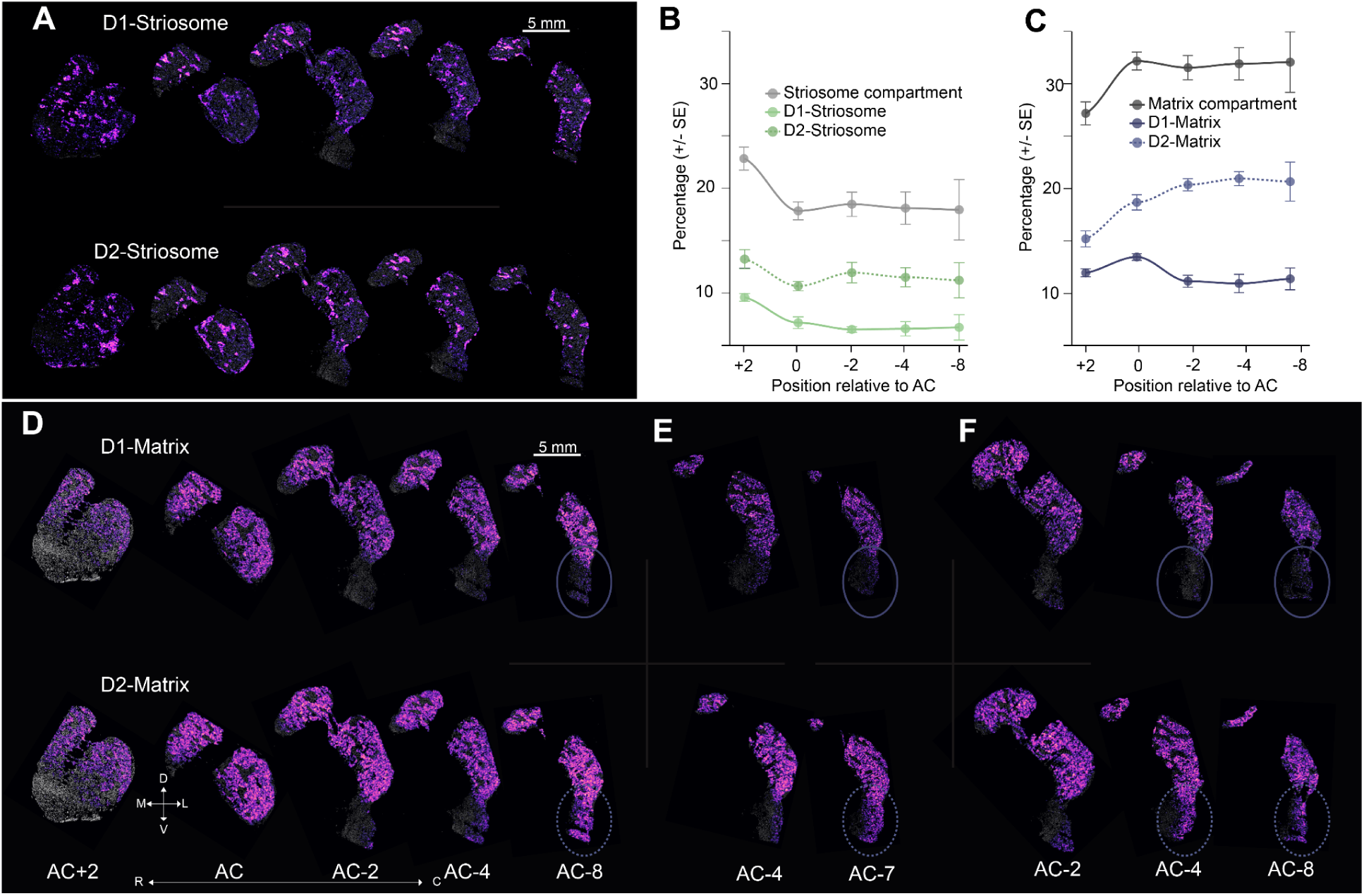
Spatial Patterning of Dorsal Striatum Medium Spiny Neurons Across Rostral-Caudal Axis. A. Cell type density map highlighting differential spatial localization of D1- and D2-Striosome across 5 levels along the RC axis (N=1). B. Regional-cell type relative abundance gradient scatter plot smoothed using LOESS across the rostral-caudal axis. For the dorsal striatum, counts of D1- and D2-Striosome—combined and separately—were aggregated across subjects and their relative proportions were calculated as percentages.Error bars represent ± SEM across sections within each region. C. Regional-cell type relative abundance gradient scatter plot smoothed using LOESS across the rostral-caudal axis. For dorsal striatum, counts of D1- and D2-Matrix—combined and separately—were aggregated across subjects and their relative proportions were calculated as percentages. D. Cell type density map highlighting differential spatial localization of D1- and D2-Matrix across 5 levels along the RC axis, highlighting an increase in D2-matrix caudally (circled area). E-F. As in D, for the caudal sections in two additional animals.

### MSN Subtype Specific Polygenic Disease Associations

Previous studies have used postmortem human and/or mouse single cell data in combination with GWAS results to demonstrate that genomic risks are localized to cell types.^37–41^ Our NHP single cell data has some potential advantages. First, we controlled the perfusion and tissue collection, and there was very little time for postmortem transcriptional or epigenomic activity, as can occur with long or variable postmortem intervals common in human samples.^42^ Second, the phylogenetic proximity between Old-World monkeys and humans is far greater than between humans and rodents. Third, our data allowed us to distinguish novel MSN subtypes. Therefore, we used our high-subtype-resolution rhesus macaque single cell data to examine, independently, epigenomic and transcriptional predictors of human diseases.

We leveraged 121 GWAS datasets encompassing psycho-social traits, neurodegenerative diseases, psychiatric disorders, substance abuse disorders, pain, sleep and metabolic traits (Supplementary table S2). Genetic variants associated with these traits were mapped to cell type-specific open chromatin regions (OCRs) identified by ATAC-seq, using the stratified linkage disequilibrium regression score (S-LDSC).^43,44^ To validate our approach, we first examined major cell classes with known disease associations (Supplementary table S4).^9,45,46^ This analysis revealed that microglia were associated with Alzheimer’s and age-related macular degeneration (AMD), neurons were related to schizophrenia, and MSNs specifically were related to substance use disorders (Supplementary Figure 5A). The concordance between our findings and classic functional studies of the brain supports the validity of our approach.

We next applied S-LDSC to the major MSN subtypes in the NHP Striatum (Supplementary table S5). The first thing we noted was that each subtype was implicated in a wide variety of diseases. Therefore, we normalized significant relationships between MSN subtypes and disease predictors and highlighted those relationships with a z-score > 1 (Figure 5A-E, “+”) (Methods). Regarding the novel RGS6-positive subtypes, both D1-VS-RGS6 and D2-VS-RGS6 subtypes exhibited strong enrichment for psycho-social traits related to intelligence ^47^ and education attainment ^48^ (Figure 5A,B). However, D2-VS-RGS6 was more strongly associated with psychiatric disorders such as schizophrenia and bipolar disorder, compared to D1-VS-RGS6 and all other MSN subtypes (Figure 5B,G). These results demonstrate unexpected disease relationships for the novel VS subtypes, suggesting that the entire striatum, rather than only the traditionally identified associative striatum, is implicated in crucial neurological traits and neuropsychiatric diseases.

**Figure 5.**
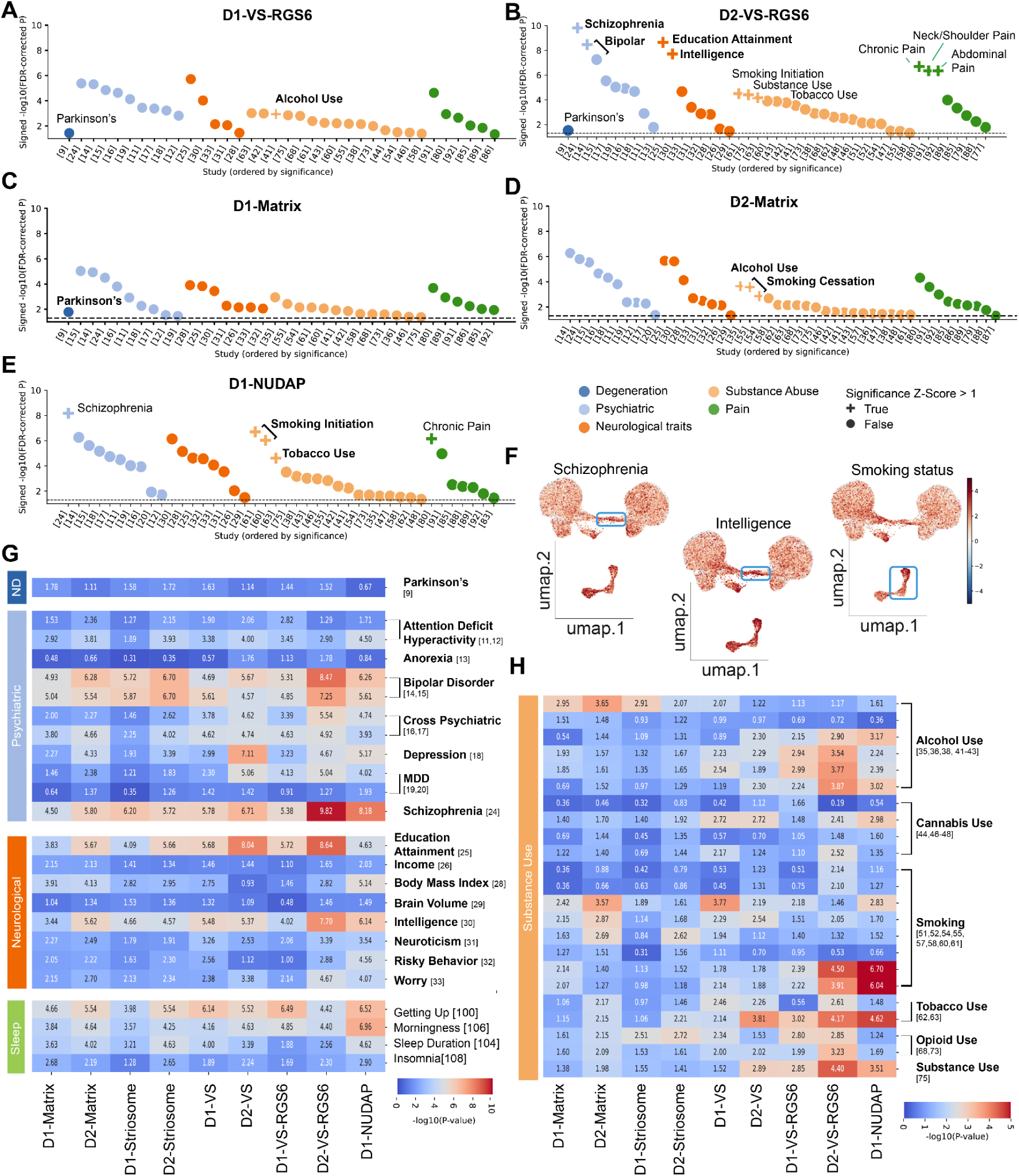
Dissecting Polygenic Disease Risk Across Cell Types in the Human Striatum. A-E. Scatterplot of s-LDSC heritability enrichment for D1-VS-RGS6, D2-VS-RGS6, D1-Matrix, D2-Matrix and D1-NUDAP paired with the significantly associated phenotypes. x-axis: GWAS studies and y-axis: signed -log10 adjusted p-value. FDR values were z-scored across cell types and GWAS studies. A “+” sign is assigned to a cell type–trait pair when both the within-study and within-cell-type z-scores exceed one standard deviation above the mean (z > 1). F. Mapping scDRS polygenic enrichment score in individual cells on low dimensionality projection of MSN subtypes (UMAP). G-H. Heatmap for significance of s-LDSC heritability enrichment across MSN subtypes-Disease pairs. P-values are FDR-adjusted across 11 subtypes and 121 traits, log transformed and multiplied by the sign of the coefficient of association (color scale bar: signed -log10 adjusted p-value). Relevant traits pertaining to neurodegenerative, psychiatric, neurological, sleep and substance use (Table S2).

Regarding the previously described subtypes, D1-matrix MSNs were more strongly related to PD than D2-matrix MSNs (Figure 5C). In contrast, psychiatric risk predictors and predictors of neurological traits, such as intelligence and educational attainment, were more strongly enriched in D2-Matrix MSNs compared to D1-Matrix MSNs (Figure 5D). Interestingly, given the enrichment of D2-Matrix MSNs in the caudal PUTv, this cell type was also strongly associated with substance use disorders (Figure 5D). D1-NUDAPs, compared to all other subtypes, were most strongly associated with substance use disorders (Figure 5E, H).

We saw the same trends when we examined the MSN subtype specific transcriptional profiles, rather than the epigenomic profiles. We applied single-cell disease relevance score (scDRS) ^49^ to examine genetic signatures for 24 human neurological diseases and traits (Supplementary table S2, S3). We found genetic predictors for neurological traits such as intelligence and educational attainment, as well as psychiatric diseases such as schizophrenia and bipolar disorder, were enriched in D2-VS-RGS6 and other VS subtypes (Figure 5F, S5B). This concurrence between the epigenomic and transcriptomic results firmly demonstrate MSN subtype specific roles in health and disease.

## DISCUSSION

Here we combined three cutting edge molecular neuroscience techniques - snRNA-Seq, snATAC-Seq, and spatial transcriptomics - to provide a comprehensive account of MSN subtypes and their spatial distributions in Old-World monkey striatum. We used multi-omic single cell sequencing to gain transcriptomic and open chromatin profiles from MSNs across four anatomically and functionally distinct regions. We used the transcriptomic profiles to identify 11 MSN or MSN-like subtypes, including two VS subtypes that were not previously described (Figure 1). We showed that these novel cell types, D1-VS-RGS6 and D2-VS-RGS6, were distinct from other MSN subtypes, but retained the limbic characteristics of subtypes previously described in the NAc (Figure 2). Both novel subtypes inhabited distinct spatial niches. The D1-VS-RGS6 subtype was preferentially located in traditionally limbic territories, including the medial NAc and in the ventro-medial CDh. In contrast, the D2-VS-RGS6 subtype was most abundant in PUTv and appeared to constitute an interface with the amygdalo-striatal transition zone. The presence of these subtypes across the RC span of the VS demonstrates the fundamentally limbic characteristic of the VS regions located outside of and caudal to the NAc (Figure 3). In the DS, we found that D1- and D2-striosome MSNs were over-represented rostral to the AC, compared to all other examined levels. In contrast, the matrix compartment MSNs increased, as a percentage of all DS cell types, in the caudal direction. This increase was driven by D2-Matrix MSNs, which populated the PUTv beyond the caudal extent of the amygdala (Figure 4). Finally, we demonstrated that genetic predictors of neurological traits and disease risks are preferentially localized to specific MSN subtypes (Figure 5). Overall, this study provides a systematic framework for understanding and further investigating the diverse functions and dysfunctions of the primate striatum.

MSNs are the most common neuron type in the striatum.^35^ They receive most striatal inputs and, as far as we know, they are the sole output neurons. Thus, they are crucial for proper basal ganglia function. MSNs are commonly considered in the context of two factors, (1) the ‘direct’ and ‘indirect’ pathways, and (2) the D1 and D2 dopamine receptor subtypes. The direct pathway projects directly to the internal segment of the Globus Pallidus (GPi) and the *substantia nigra pars reticulata* (SNpr), whereas the indirect pathway projects indirectly to the GPi and SNpr via the external segment of the Globus Pallidus (GPe) and subthalamic nucleus (STN).^12^ Landmark research demonstrated that lesions to dopamine neurons caused different gene expression changes in direct and indirect pathway MSNs^50^ and led to the hypothesis that hyperkinetic and bradykinetic symptoms observed in Huntington’s and Parkinson’s disease, respectively, were pathway specific.^51^ The cell type specific source of this difference was revealed with the discovery that D1 was expressed only in MSNs that form the direct pathway projection to GPi and SNpr, whereas D2 was expressed only in MSNs that form the indirect pathway projection to the GPi and SNpr.^12–14^ This molecular-functional distinction has been crucial for understanding the striatum’s central role in PD, which is marked by motor symptoms arising from the selective loss of dopaminergic neurons in the substantia nigra pars compacta (SNpc) projecting to the dorsal striatum. This dopaminergic degeneration results in repression of the direct pathway, shifting the balance towards the indirect pathway and ultimately resulting in movement inhibition.^52,53^ Leveraging this cell type specific circuit architecture, new gene-based therapy circuit modulation tools are being developed to selectively manipulate PD-affected circuitry. For example, using promoter-driven retrograde adeno-associated virus (AAV) and a chemogenetic effector for selective targeting and activation of DRD1-expressing MSNs, the motor signs were rescued in a primate model of PD.^54^ The single cell techniques we demonstrate here have the potential to take us deeper, both with regard to cell type specific neuroanatomy and with the development of targeted therapies to ameliorate the symptoms of PD and the numerous other disorders associated with circuit dysfunction in the BG.

With these goals in mind, we applied single cell multi-omics to demonstrate that there are 11 distinct MSN subtypes in the NHP striatum. Two of these, D1-VS-RGS6 and D2-VS-RGS6 were previously unknown, but expressed known markers of VS MSNs, including *GRIA4*.^26^ We also uncovered a novel VS marker gene: *FGFR2*, that was enriched in all four major VS cell types. Despite their name suggesting fibroblast specificity, fibroblast growth factor receptors have been shown to play key roles in modulating synaptic plasticity in D2-receptor expressing neurons.^55^ Although the D1-VS-RGS6 and D2-VS-RGS6 neurons exhibited shared gene expression profiles with previously described VS cell types, differential gene expression and chromatin accessibility provided compelling evidence that they were distinct cell types. In fact, there was greater separation between RGS6-positive and previously described VS MSNs than between canonical D1- and D2-Matrix MSNs (Figure 2C, D). Thus, we revealed two novel VS cell types characterized by the expression of *RGS6*.

RGS proteins are a large family of GTPase-activating proteins that play a critical role in neurotransmission by regulating the duration and magnitude of G protein coupled receptor signaling.^36^ Surprisingly, across this whole family, only *RGS6* was enriched in specific MSN subtypes (Figure 2E), suggesting that *RGS6* has a critical role in cell type specific striatal functions. In dopamine neurons, *RGS6* inactivates the D2-auto-receptor and has been suggested to be neuroprotective through promoting their survival and rescuing PD-associated motor deficits.^56,57^ Interestingly, loss of *RGS6* expression was selective to dopaminergic neurons in the ventral SNpc in a mouse model of PD.^56,58^ *RGS6* expression in MSNs has not been previously described, but *RGS9* inhibition of D2-receptor signaling facilitates excitability of MSNs^57^ and appears to be neuroprotective in rodent models of PD.^59,60^ Therefore, it is critical for future studies to examine the RGS6-positive subtypes in the context of PD.

Our data indicate that the VS contains four major MSN subtypes, the two RGS6-positive subtypes and the two VS subtypes revealed by our prior analysis of MSNs in the NHP striatum.^26^ Those experiments focused on the region around and rostral to the AC, and the spatial analysis was limited to single molecule Fluorescent In-Situ Hybridization (FISH).^26^ Those data indicated that the transcriptionally-distinguishable VS subtypes were restricted to the NAc shell. Therefore, we had denoted these as D1- and D2-Shell MSNs. The current data, with broader sampling of single cell multi-omics and cutting-edge spatial transcriptomics, revealed that these ‘shell’ subtypes were found within and beyond the NAc shell, and even beyond the NAc. Therefore, we relabeled them as D1- and D2-VS. The four limbic MSN subtypes were abundant at rostral RC levels that included the NAc, but they were also abundant in caudal sections, where they were found in the medial CDh, PUTv, and CDt, striatal regions that receive inputs from the amygdala.^32^ This finding suggests that the PUTv is a caudal extension of the limbic striatum, rather than a ventral extension of sensorimotor territories.

The spatial transcriptomics also revealed rostral-caudal shifts in the proportions of matrix-striosome compartments, where matrix cell types, particularly D2-matrix, were more abundant in the caudal striatum relative to strisomes. This finding is aligned with previous reports indicating loss of compartmentalization in the ventral half of the caudal striatum.^61^ However, our data extend these observations by characterizing this loss as a redistribution of striatal compartments, partially driven by the spatial gradient of *DRD1* and *DRD2* expression along the R-C axis.

The presence of limbic-type MSNs in the PUTv aligns broadly with prior neurochemical evidence for limbic continuity between rostral and caudal portions of the primate VS.^32^ That study revealed a calbindin-poor region adjacent to the amygdalostriatal region that was interpreted as an extension of the NAc shell, whereas the more lateral PUTv was calbindin rich and considered an extension of the traditional striatum. However, the cell types that we observed are not consistent with the conclusion that PUTv is an extension of the NAc shell. Our data indicate that the *CALB1* poor zone in the caudal striatum was populated by MSN subtypes that are different from those located in the NAc shell. Instead, this region contained the D1-VS-RGS6 and D2-VS-RGS6 subtypes, whereas the *CALB1* rich lateral PUTv contained the D1-VS and D2-VS subtypes. These findings highlight the limitations of single-marker definitions and emphasize the power of single-cell, multimodal analyses for refining our understanding of striatal organization. They also suggest that PUTv is a highly complex area requiring more investigation.

Corticostriatal inputs to PUTv and the adjacent CDt arise from the visuotemporal areas, including IT cortex.^62,63^ Single cell recordings show that MSNs in this territory code for the learned value of objects.^64^ Our spatial transcriptomics showed that, at the level of the amygdalostriatal transition zone, the four major VS cell types occupied distinct medial-lateral territories, with D2-VS-RGS6 the most medial and D1-VS the most lateral. As we continued beyond the caudal extent of the amygdala, around 8 mm beyond the AC, there emerged an abundance of D2-Matrix-MSNs, but a striking paucity of D1-Matrix MSNs (Figure 4D-F). This differential enrichment of matrix compartment MSNs in PUTv seems to explain the relative increase in D2- to D1-Matrix MSNs in the caudal striatum (Figure 4C), but also raises important questions about circuit organization. Tract tracing indicates that cells in this region contribute to the direct and indirect pathways.^65^ Given the relative scarcity of D1-Matrix MSNs, what cell types give rise to the direct pathway projections to the SNpr? Addressing this question will require new experimental approaches. The methods and data presented here suggest a path forward: spatial transcriptomic techniques, when combined with conventional and polysynaptic tracing tools, can elucidate how specific MSN subtypes relate to their afferent and efferent connectivity. In parallel, our snATAC-Seq data identify cell type specific enhancers that can be incorporated into AAV vectors, providing the foundation for causal investigations linking defined MSN subtypes to circuit function and behavior. ^66^

One of the most crucial future directions is revealing the cell type specific bases for disease mechanisms. As we highlighted in the introduction, the basal ganglia are implicated in a wide variety of diseases, including movement disorders,^7^ psychiatric diseases,^8^ and substance use disorders.^9^ In order to facilitate the search for cell type specific bases for disease mechanisms, we applied two computational methods, S-LDSC and scDRS, to examine cell type specific enrichment of disease and trait-predicting genomic elements. Because many of these elements are located in non-coding regions of the genome,^67^ we focused mainly on S-LDSC because it examines these noncoding regions in the snATAC-Seq data. We validated the performance of S-LDSC on our NHP single cell dataset by looking at well known relationships between disease predictors and broad cell classes. This validation was successful: Alzheimer’s disease (AD) risk variants were exclusively enriched in microglia,^18,68,69^ consistent with extensive evidence implicating microglia to be key cell mediators of neuroinflammation in the AD brain.^45^ Microglia were also the only cell type to be significantly associated with the genetic risk for AMD, reinforcing shared molecular mechanisms between AD and AMD.^70^ Schizophrenia risk variants were most enriched in neurons,^40,71,72^ whereas substance use disorders were most strongly associated with a neuron subclass, MSNs. Given this strong validation against known disease relationships, we applied S-LDSC to the MSN subtypes and revealed some surprising associations. Genetic predictors of intelligence and educational attainment were preferentially localized to both D1-VS-RGS6 and D2-VS-RGS6 subtypes, whereas, intriguingly, schizophrenia and bipolar disorder were most strongly localized to D2-VS-RGS6. This is intriguing because this cell type is not located in the associative regions implicated in these psychiatric disorders. The localization of PD risk factors to the RGS6-positive subtypes was not as strong, but survived strict multiple comparison correction. This strongly implicates these subtypes in PD, despite the fact that they are located outside of the sensorimotor striatum that influences movements.^2,3^ Genetic risk predictors for substance use disorders were most strongly associated with D1-NUDAPs, which is of potential interest due to their high expression of μ-opoid receptors. These gene associations can provide the starting point for causal studies to link diseases with cell type- and circuit-based mechanisms.

Altogether, the current results provide a comprehensive picture of genetic variability in MSNs across the NHP striatum and indicate how that genetic variability between MSN subtypes might be related to disease. More importantly, the spatial transcriptomics results provide a roadmap for linking cell types with large-scale brain networks, and the multi-omic database contains thousands of cell type specific open-chromatin regions that might contain the key to delivering circuit-targeted experimental and therapeutic genetic payloads. Thus, the current results set the stage for future studies to reveal cell type specific striatal functions.

## SUPPLEMENTARY MATERIALS

Supplementary Figures 1-5

Supplementary Tables 1-5

Methods

Acknowledgments

---

**Supplementary Figure 1.**
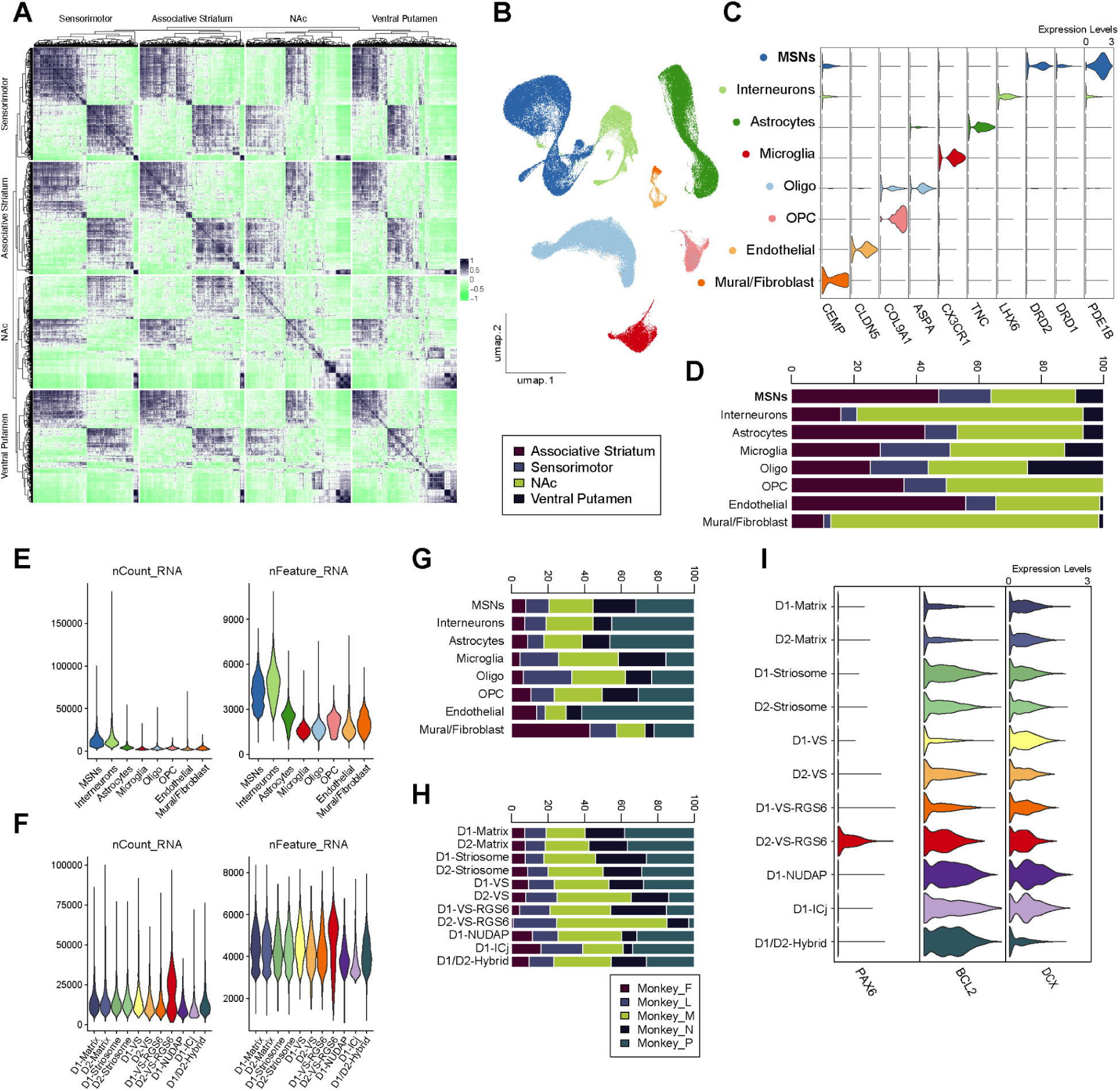
Cell Type Distribution Across Subjects-Regions and Quality Control Metrics For snRNA-seq data. A. Hierarchically clustered heatmap of the gene expression vectors cosine similarity between each pair of cells, rows and columns are split by striatal regions (in A), highlighting transcriptional regional heterogeneity. snRNA-seq dataset was down-sampled to 2000 cells comprising 500 randomly sampled cells per each of the four regions. Cosine similarity was computed between embedding vectors (30 PCs, principal components) per cell. B. Low dimensionality projection of striatal cell classes (UMAP), after QC filtering, integration of striatal neuronal and non-neuronal nuclei from 4 striatal regions and 5 rhesus macaques, clustering and major cell classes annotations. C. Violin plots for the major cell classes-specific marker genes neuronal and non-neuronal striatal nuclei. Normalized gene expression per cell type is averaged across regions and animals. D. Stacked barplot showing the relative regional proportions per cell class for the four functionally defined striatal regions. E-F. Averaged per-nuclei quality control (QC) metrics across major cell classes (E) and MSNs subtypes (F). G-H. Stacked barplot showing relative proportions of cells from each of the 5 rhesus macaques per cell class (G) and subclass (H). Cell types are represented across all animals and are not driven by a single subject. I. Neural progenitor cell markers, *DCX*, *BCL2* and *PAX6* gene expression for the 11 cell subtypes of MSNs. Normalized gene expression per cell type is averaged across regions and animals.

**Supplementary Figure 2.**
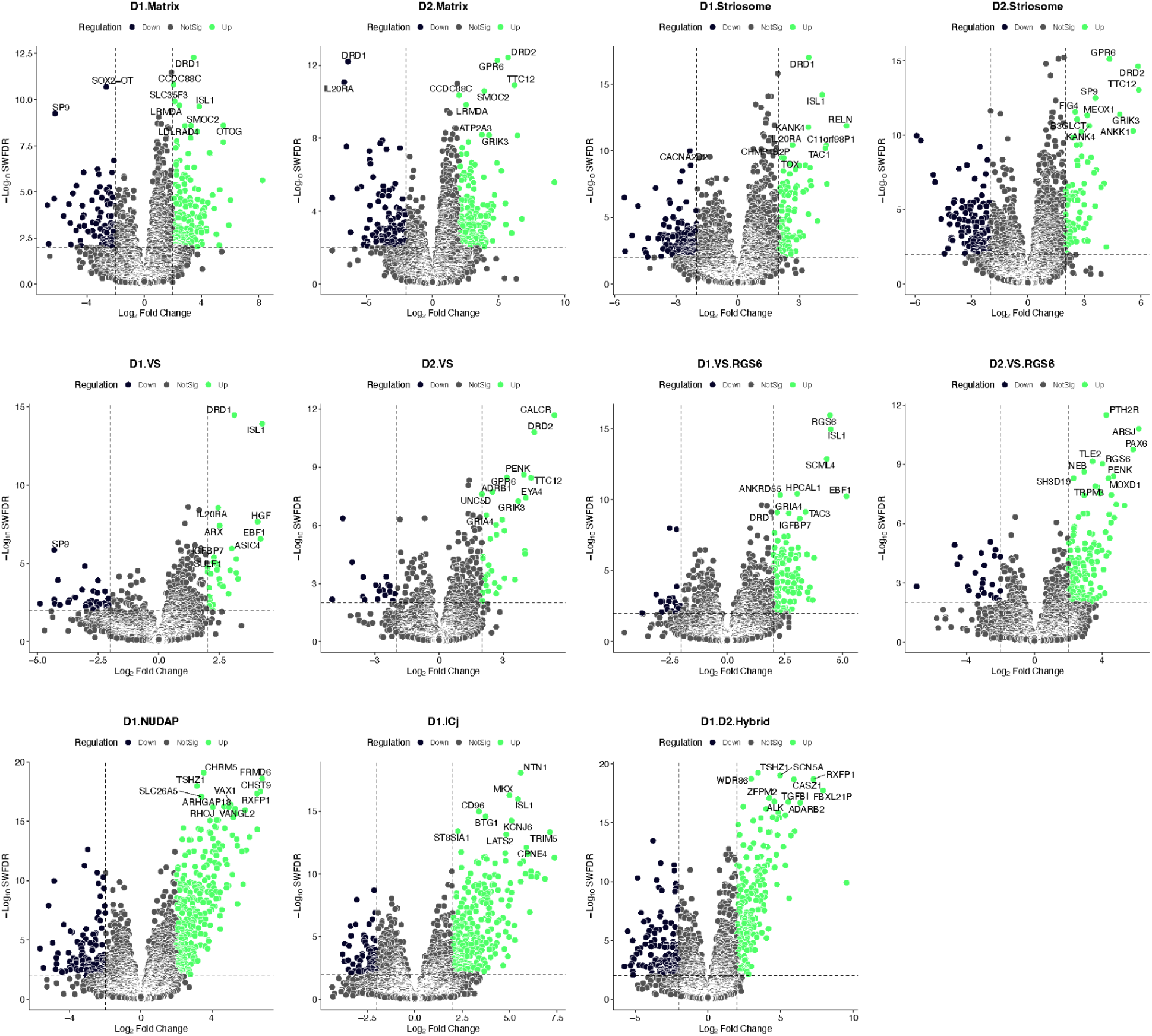
Differential Gene Expression Profiles for MSNs Cell Types. Differential gene expression analysis for each of the eleven identified MSN subtypes in contrast to all other MSN subtypes using Limma-Voom (Supplementary table S1). In each volcano plot, x-axis: Log2(Fold Change), y-axis: negative Log10(Weighted FDR) and the top 10 genes for each cell type are annotated.

**Supplementary Figure 3.**
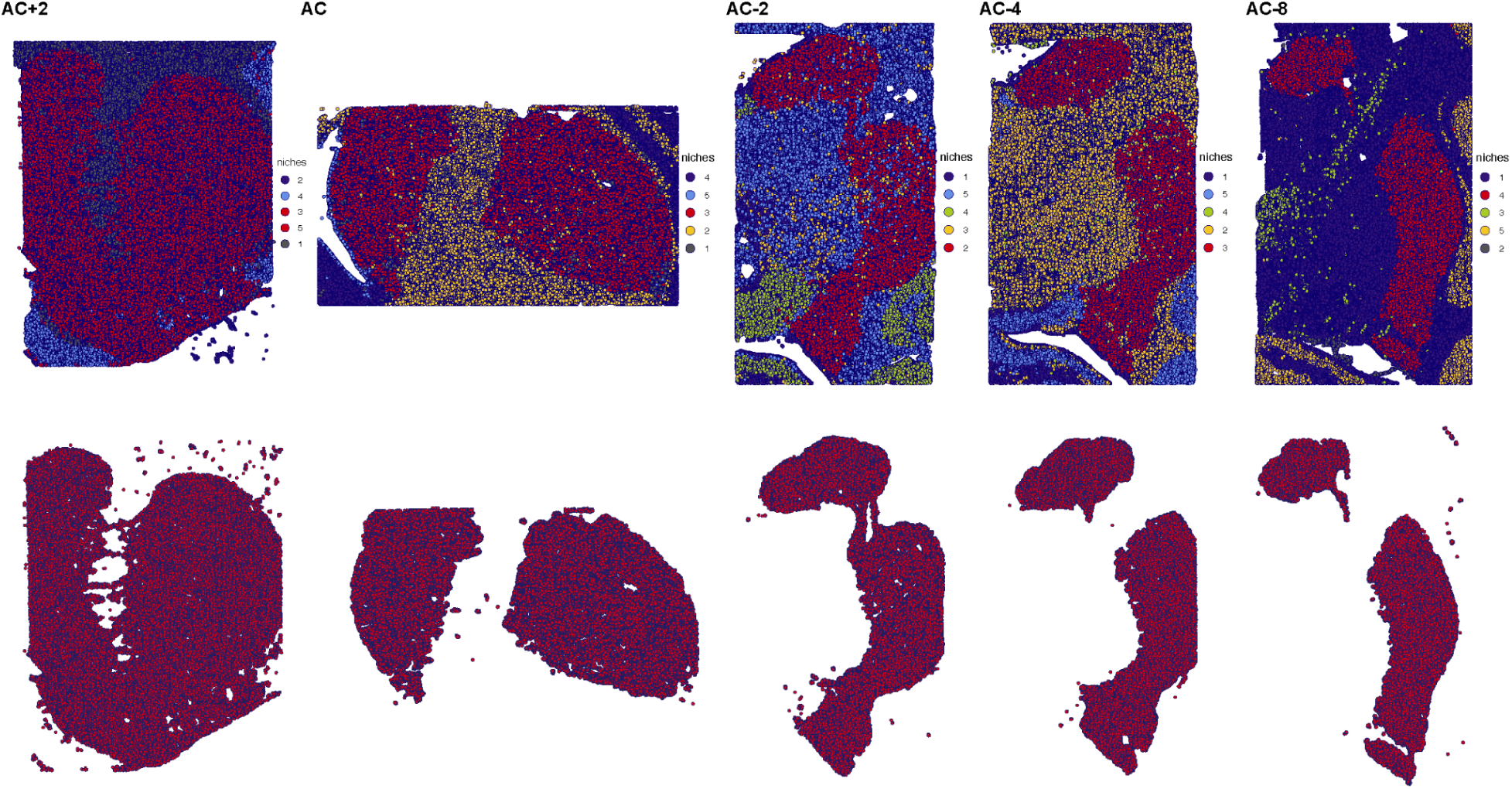
Spatial Transcriptomics Striatal Tissue Dissection. Spatial niche analysis for midbrain tissue sections used for xenium in-situ expression assay across the R-C axis using the anterior Commissure (AC) as the reference level (AC+2, AC, AC-2, AC-4, AC-8). Striatal niche is assigned red color in all sections.

**Supplementary Figure 4.**
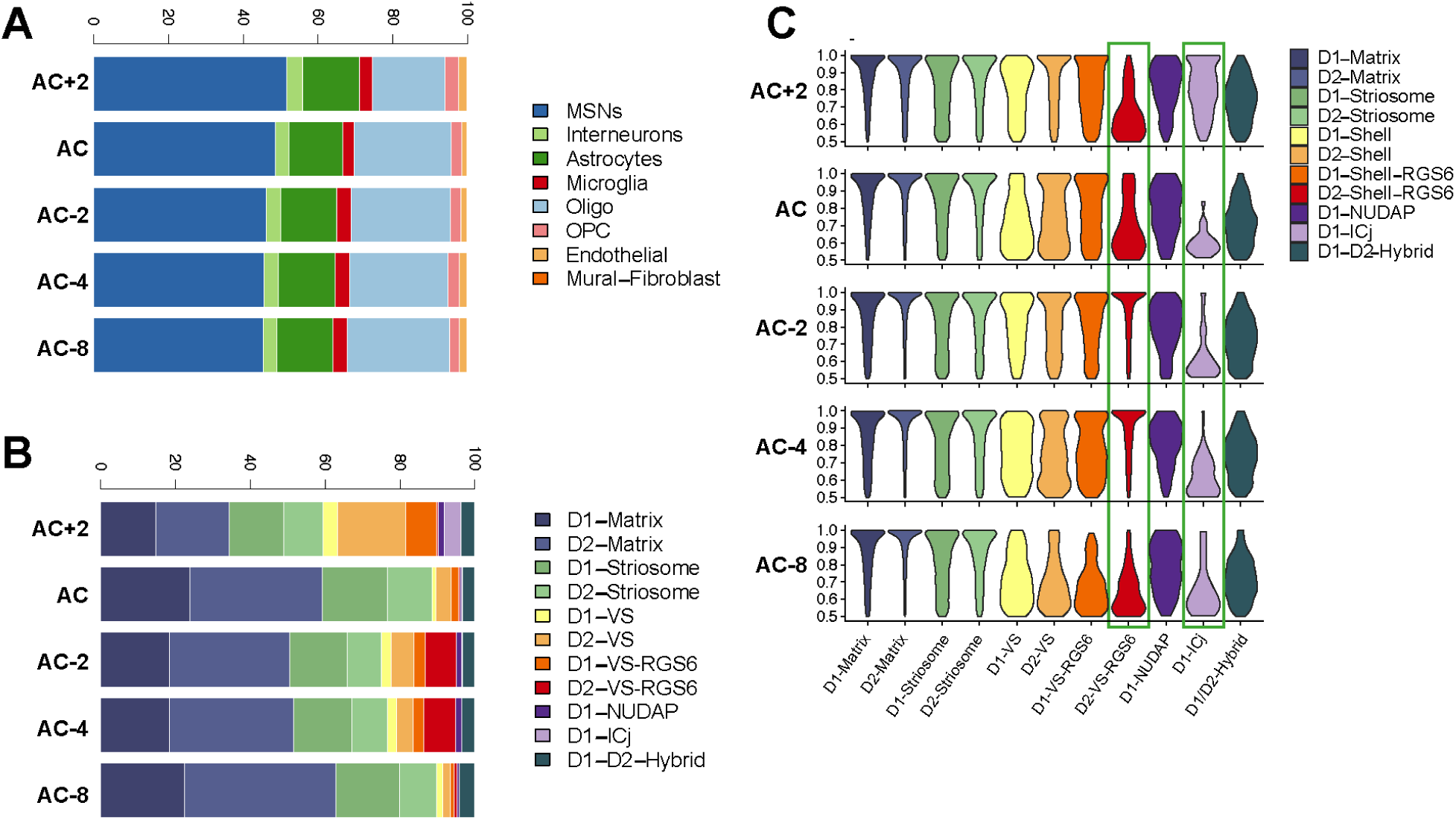
Regional Cell Type Prediction and Composition for Spatial Transcriptomics. A-B. Stacked barplot showing regional-cell type relative proportions of major cell types (A) and MSNs subtypes (B) in five sections across the RC axis. C. Violin plots of the prediction scores for MSNs subtypes (thresholded at 0.5). Green rectangles highlight the increase in prediction scores depending on the abundance of a cell type, D2-VS-RGS6 in caudal sections and D1-ICj in rostral section.

**Supplementary Figure 5.**
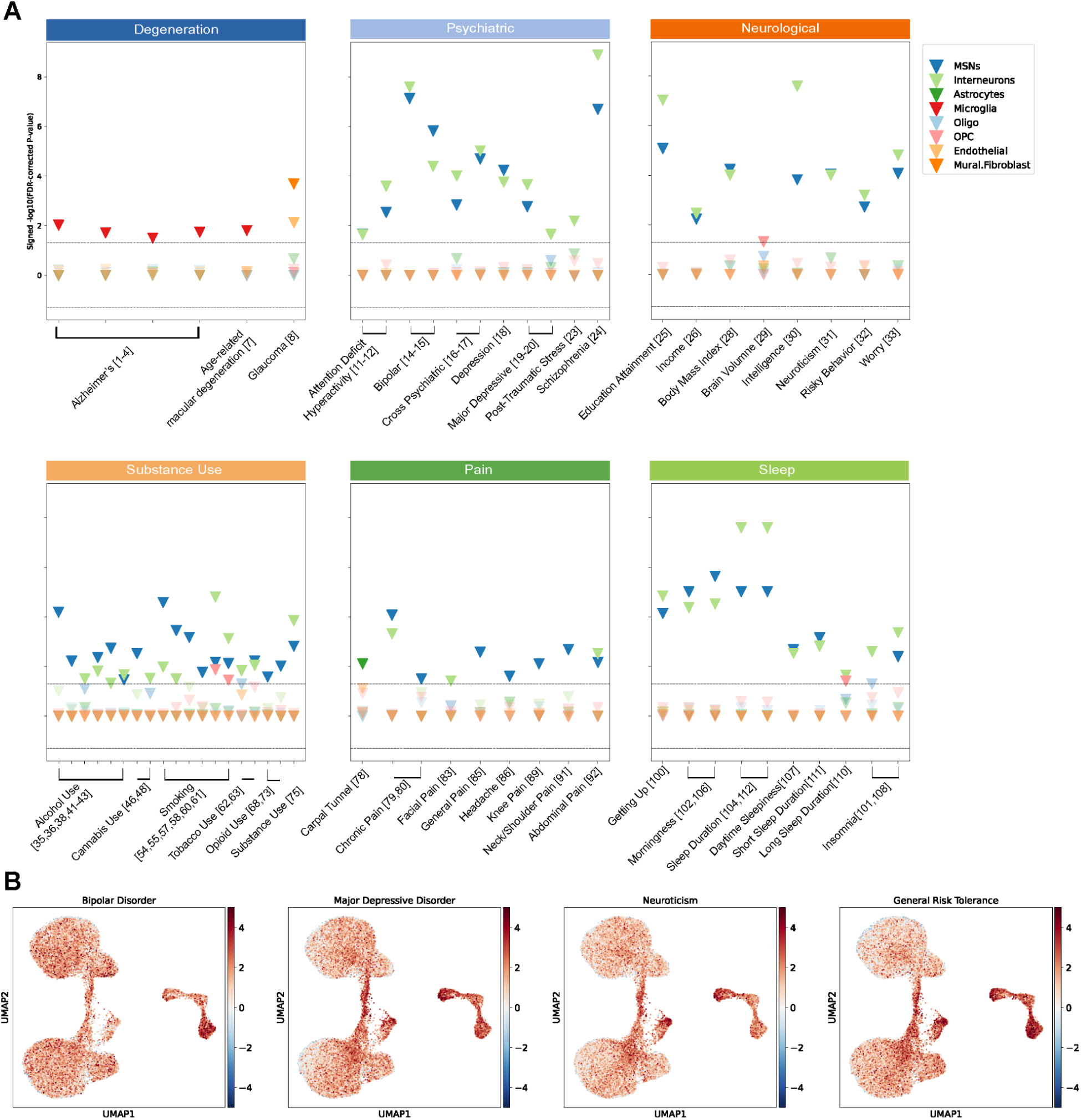
Dissecting Polygenic Disease Risk Across Cell Types in the Primate Striatum. A. Scatterplot for significance of LDSC heritability enrichment across striatal major cell type-Disease pairs. P-values are FDR-adjusted across 11 subtypes and 121 traits, log transformed and multiplied by the sign of the coefficient of association (color scale bar: signed -log10 adjusted p-value). Relevant traits pertaining to neurodegenerative, psychiatric, neurological, sleep and substance use (Table S2). Black dotted line marks FDR=0.05. Bold triangles represent significant associations, while faded triangles represent non-significant associations. B. Mapping scDRS polygenic enrichment score in individual cells on low dimensionality projection of MSN subtypes (UMAP).

## EXPERIMENTAL MODEL

### Non-Human Primates (NHPs)

All animal procedures were in accordance with the National Institutes of Health Guide for the Care and Use of Laboratory Animals and approved by either the University of Pittsburgh’s or the Emory National Research Primate Center’s Institutional Animal Care and Use Committee (IACUC). Rhesus and Cynomolgus Monkeys were single- or pair-housed with a 12 h-12 h light-dark cycle. Experimental procedures included tissue from 7 rhesus and 3 cynomolgus macaques with age range 3-13 years old. Rhesus macaques included: monkey SM019 (male, age at perfusion=3 years), monkey SM045 (male, age at perfusion=3 years), monkey SM020 (female, age at perfusion=3 years), SM057 (female, age at perfusion=4 years), SM013 (female, age at perfusion=5 years), monkey AG001 (female, age at perfusion=9 years) and monkey SM007 (female, age at perfusion=12 years). Cynomolgus macaques included: monkey SM074 (male, age at perfusion=5), monkey SM056 (male, age at perfusion=9) and monkey SM039 (male, age at perfusion=13).

## METHODS

### Tissue Collection and Harvesting

Tissue collection for snRNA-seq (SM007 and SM013) was performed as previously described.^26^ For multi-ome experiments, animals were anesthetized with ketamine (15 mg/kg IM), administered an overdose of pentobarbital sodium (40 mg/kg IV or IP), and perfused transcardially with approximately 4 L of oxygenated ice-cold artificial cerebrospinal fluid (ACSF).The brains were removed and cut into ∼3-4 mm thick slabs in the coronal plane. The striatal regions of interest, associative striatum, nucleus accumbens (NAc), sensorimotor putamen, and ventral putamen (PUTv) were dissected with microforceps. Striatal tissues from rhesus macaques (n=5, 3 females and 2 males, age range at perfusion 2-12 years including SM007, SM013, SM019, SM020, SM045).

For spatial transcriptomics, animals were perfused with RNase-free phosphate buffered saline (10X PBS, Thermo Fisher Scientific, Cat# AM9625 diluted 1:10 in Milli-Q water) followed by 4% paraformaldehyde (PFA, Sigma-Aldrich, Cat# P6148). Brains were extracted and post-fixed with 4% PFA and cryopreserved with sucrose gradients (10% and 20% in 4% PFA, and 30% in PBS). Brains were blocked in the coronal plane, frozen, and cut into coronal sections 20μm thick. Selected sections were mounted on Xenium slides and stored at -80°C before spatial transcriptomic analysis.

### Single-nucleus RNA Sequencing

For monkeys SM007 and SM013, samples were run using the 10x Chromium Single Cell 3′ Reagent kits, v3.1 Chemistry. Nuclei isolation was carried out as previously described.^26^ Library preparation was performed following the standard 10x protocol for v3.1 chemistry. The experimental protocol followed these steps: 1. Reverse transcription of mRNAs within Gel beads-in-emulsion (GEMs) after running through a 10x Genomics Chromium controller, 2. Breaking the emulsion with a recovery agent (10x Genomics, Cat# 220016), 3. Purification of cDNAs with Dynabeads (10x Genomics, Cat# 2000048), 4. Amplification of cDNAs and purification with SPRIselect reagent (Beckman Coulter, Cat# B23318), 5. After evaluating the cDNA quality using an Agilent Bioanalyzer 2100, libraries were prepared following fragmentation, end repair, A-tailing, adaptor ligation, and sample index PCR, 6. Quantification of the libraries by qPCR using a KAPA Library Quantification Kit (KAPA Biosystems, Cat# KK4824) and 7. Pooling libraries from individual monkeys together and loading them onto NovaSeq S4 Flow Cell Chip. We sequenced samples to the depth of 200,000 reads per nucleus.

### Single-nucleus RNA and ATAC multi-omics Sequencing

For monkeys SM019, SM020 and SM045, samples were run using the 10x Chromium Single Cell Multiome Library & Gel Bead Kit (10X Genomics, Cat# PN-1000283), providing snRNA-Seq and snATAC-Seq from the same nuclei. The snATAC-Seq and snRNA-Seq libraries were prepared according to the standard protocol. The experimental protocol included these steps: 1. Breaking the tissue into small pieces with a wide-bore pipette tip followed by a regular-bore pipette tip, then filtered through a 30 μm MACS SmartStrainer, 2. After spin down, cell lysis with 0.1X lysis buffer (10mM Tris-Hcl, 10mM NaCl, 3mM MgCl2, 0.1% Tween 20, 0.1% NP-40, 0.01% Digitonin, 1% BSA, 1mM DTT, 1U/ul RNase inhibitor), 3. After washing, nuclei resuspension with diluted nuclei buffer (10x Genomics, PN-2000153). We adjusted the nuclei concentration to 2,900-7,260 nuclei/μl, based on the targeted nuclei recovery of 9,000, 4. Nuclei transposed and ran through the 10x chromium controller, followed by reverse transcription, 5. Pooling libraries and loading them onto a NovaSeq 6000 S4 Flow Cell. We sequenced each sample to the depth of 150,000 x 150 bp paired-end reads per nuclei.

### Single-nucleus RNA Sequencing Analysis

Sequencing BCL files were converted to FASTQ file format, which is the default input for secondary data analysis. Reads were aligned using a custom rheMac10 transcriptome reference ^73^, which was created to extend macaque gene annotations with human annotations using the UCSC liftOver tool. Using the Cell Ranger analysis pipeline, reads from FASTQ files were aligned to the reference genome using the custom annotation file, which generates count matrices for each monkey. The final set of genes comprised 34,455 genes, after removing ribosomal genes. Seurat pipeline default clustering was performed prior to decontaminating data from ambient RNA using SoupX.^74^ Stringent quality control was applied to remove low quality cells using Seurat (nFeature_RNA > 2500 & nFeature_RNA < 10000 & percent.mt < 5) and doublets were removed using DoubletFinder^75^. Seurat v5 was used for downstream data processing.^76^

For the integrative analysis, caudate, putamen and nucleus accumbens (NAc) brain regions were initially processed, annotated and integrated for all monkeys (n = 5). The caudal ventral putamen sample from 3 monkeys was independently processed, then integrated firstly across monkeys and secondly with the 3-regions integrated object. Prior to integration, we performed standard log-normalization and identified variable features using Seurat’s logNormalize and FindVariableFeatures functions. Anchors were identified using FindIntegrationAnchors function with default parameters followed by integration using IntegrateData function from Seurat. This yielded an integrated expression matrix for all nuclei across 5 monkeys and 4 striatal regions. Principal component analysis (PCA) was run on the scaled and integrated object followed by uniform manifold approximation projections (UMAP) for visualization using RunPCA and RunUMAP functions. Louvain clustering was performed at a resolution that provided clear cluster separation for glial and neuronal cell types. We annotated major cell classes based on marker genes calculated using FindMarkers function. Based on previously reported MSNs (MSNs) marker genes, PPP1R1B, BCL11B, and PDE1B ^26^, MSNs clusters were defined and glia and interneurons were removed. Variable features were re-identified followed by re-calculating principal components, re-running UMAP dimensionality reduction and re-clustering. For snRNA-seq, integration of all nuclei across 4-regions and 5 monkeys yielded a total of **158,728 nuclei** with 1,500 variable features. Integration of MSNs across 4-regions yielded **39,352 MSNs nuclei** with 1,500 variable features. Resolution was adjusted initially to low-resolution, 0.1, to provide a coarse representation of the clusters. Resolution was adjusted to high-resolution, 0.8, to provide a fine representation of clusters for downstream analyses.

### Single-nucleus RNA Sequencing Differential Gene Expression Analysis

Using the Limma-Voom approach ^77,78^, we performed a differential gene expression analysis to identify cell type specific transcriptional signatures while accounting for sample-level variation and unmodeled confounders ^79^. This analysis consisted of the following steps: **(1) Pseudobulk aggregation:** we constructed cell type x animal pseudo-bulk gene expression profiles from single cells to lower the rate of false discovery for gene-level differences usually caused by repeated measurements from single cells from the same individual. For each cell type x animal combination, raw counts were summed using aggregate.Matrix() function. Pseudobulk aggregates with less than 15 cells were excluded. Additionally, the gene detection rate (fraction of detected genes per pseudobulk) per aggregate was computed for quality control. **(2) Normalization and Variance Modeling:** raw counts were normalized using calcNormFactors() function and genes with low counts, as well as ribosomal and mitochondrial genes were excluded from the analysis. For gene expression values, we estimated observation-level precision weights modeling the mean-variance relationship and sample-specific quality weights using voomWithQualityWeights() function. To account for repeated measurements from the same animal, we computed intra-subject correlation using duplicateCorrelation() function, then recomputed expression weights incorporating sample correlations. **(3) Model Design:** we constructed two models, design matrix model *(∼ 0 + celltype + numCells + gene detection rate)* – capturing cell type effects on gene expression while controlling for technical covariates (number of cells and gene detection rate per pseudobulk sample) and a null model *(∼ 0 + numCells + gene detection rate)* – capturing covariate effects only for surrogate variables estimation. **(4) Surrogate Variable Analysis:** to capture unmodeled sources of variation, we used the full design model and null model to identify surrogate variables while preserving biological variation using the sva package. The model was adjusted by adding the surrogate variables, followed by re-estimating the expression weights and sample-level correlations. **(5) Linear Modeling and Contrasting:** we fitted a linear model with lmFit() and applied empirical Bayes moderation to stabilize variance estimates with eBayes(). For each cell type, differential expression was computed using a one-versus-all contrast design. We re-fitted the model with the contrasts using contrasts.fit() and applied eBayes() again. **(6) Covariate-Adaptive False Discovery Weight:** for the differentially expressed gene (DEG) lists, we applied multiple hypothesis correction using the FDR method while accounting for average gene expression as a covariate using lm_qvalue() from swfdr package ^80^. This approach models how p-values depend on the average gene expression, giving higher weight to informative genes and thus, detecting true DEGs. We applied FDR weighted correction within each cell type (adj.P.Val.Within) and across all cell types (adj.P.Val.Between), which is more conservative and reported in text and figures (Supplementary table S1).

### Single-nucleus Multi-ome ATAC Sequencing Preprocessing

Sequencing BCL files were converted to FASTQ file format using cellranger-arc mkfastq. snATAC-Seq reads were aligned to rheMac10 genome using fast mapper Chromap ^73,81^. This was followed by creating arrow files using the ArchR package. For gene and genome annotations, we used a custom rheMac10 annotation by mapping GRCh38.p13 human RefSeq gene annotations onto the rheMac10 genome with the liftOff tool ^26,82^. This allowed us to use ATAC-seq signal proximal to the TSS to estimate gene activity for more genes than what exists in the base rheMac10 gene annotations using the ArchR package ^83^. For downstream analyses, we implemented the pipeline previously described for preprocessing ^66^. For quality control, we performed two rounds of low quality cell filtering. The first round included fragment count filtering and doublet removal, where we discarded cells with number of fragments less than 10^3.5 and/or high probability of being doublets using filterDoublets function (cutEnrich = 0.5, cutScore = -log10(.05), filterRatio = 1). The second round involved an outlier cell scoring, where we fit a negative binomial regression to model fragment count as a function of TSS enrichment, promoter ratio and doublet enrichment of the form: *glm.nb(nFrags ∼ TSSEnrichment + PromoterRatio + DoubletEnrichment)* ^84^. The residuals from fit were standardized and nuclei with residuals more than two standard deviations from the fit (residual cutoff > 2) were excluded. For snATAC-seq, there are 66809 high quality total nuclei across 3 animals, comprising 17467 MSNs.

### Single-nucleus Multi-ome ATAC Sequencing Downstream Processing

We performed iterative latent-semantic index (LSI)-based dimensionality reduction and harmony batch correction^85^. Uniform manifold approximation and projection (UMAP) embeddings and cluster annotations were added. Cell type annotations were transferred from the corresponding snRNA-seq multi-ome dataset based on shared unique cell barcodes. Next, we identified chromatin accessibility profiles for major cell classes and MSNs subtypes separately. This process involved: (1) generating of cell type x sample pseudo-bulk replicates using addGroupCoverages function (minCells = 40, maxCells = 2000, minReplicates = 2, maxReplicates = 12), and (2) calling peaks with MAC2^86^ and constructing a union reproducible peak set using addReproduciblePeakSet function (reproducibility = “(n+1)/2”, genomeSize = 2.7e9, peaksPerCell = 500, maxPeaks = 300000, minCells = 25, cutOff = 0.1, promoterRegion = c(5000, 100)). This yielded a cell x peak matrix through ArchR. Using the peak matrix, we identified differential OCRs in each cell type relative to all other cell types as well as for each pairwise cell type comparison. Differential accessibility analysis was performed using getMarkerFeatures function (bias = c(“TSSEnrichment”, “log10(nFrags)”), testMethod = “wilcoxon”), followed by filtering using getMarkers function (cutOff = “FDR <= 0.05 & Log2FC >= 0.25”).

### LDSC-based Heritability Enrichment

We assessed associations of the regulatory profiles of striatal cell types with human genetic risk profiles for a wide spectrum of diseases and traits. We leveraged 121 GWAS datasets encompassing psycho-social related characteristics, neurodegenerative diseases, psychiatric disorders, substance abuse disorders, pain, sleep and metabolic-associated traits (Table S1). Stratified linkage disequilibrium score regression (S-LDSC) was used to estimate cell-type-specific heritability enrichment by regressing GWAS χ² statistics against LD Scores within cell-type-specific genomic annotations. ^43,44^ Conditional heritability enrichment was estimated for each disease-cell type pair, where a significant positive association represents a positive contribution of the foreground annotation to per-SNP heritability conditional on background and baseline annotations.^87^ The power of S-LDSC depends upon the annotation size and power of the GWAS. Annotations must contain >0.5% of SNPs or ∼1% of the genome to control type I error.^88^ The ∼1% of the genome thresholds was met for striatal cell types annotations. The pipeline included: (1) Call reproducible peaks using ArchR for major cell classes including medium spiny neurons (MSNs), interneurons glia, endothelial cells and mural/fibroblasts and independently within MSNs including the eleven identified subtypes. The output is foreground genomic regions of interest for each cell type and background genomic regions including a merged peak set of all striatal cell types, (2) Lift ATAC-seq peaks from rheMac10 to rheMac8 genome assembly to match cactus hal file^89^, (3) Map macaque peaks to human orthologous regions using HALPER^90^, (4) Annotate macaque-to-human lifted-over ATAC-seq peaks with LD scores using Hg38 1000 Genomes European Phase 3 European super-population (1000G EUR) cohort, (5) Prepare GWAS summary statistics for LDSC regression, datasets were processed using LDSC munge_sumstats function, and filtered to SNPs in the HapMap3 reference panel, excluding the MHC region and SNPs in regions of high inter-chromosomal LD, (6) S-LDSC based heritability enrichment per cell type annotation was estimated using the foreground annotation as the cell type-specific open chromatin regions (OCRs) relative to the background annotation, (7) Significance of association (coefficient p-value estimates) were corrected with Benjamini-Hochberg method (BH/FDR) across cell types and 121 traits (8 cell types x 121 GWAS for major cell classes and 11 cell types x 121 GWAS for MSNs subtypes), log transformed and multiplied by the sign of the coefficient of association (signed -log10 adjusted p-value). To identify MSN subtypes significantly enriched for specific traits, we first z-scored FDR values for each study across all cell types. We then binned results by cell type and major trait category and again z-scored significant FDR values within each bin. A “+” sign was assigned to a cell type–trait pair when both the *within-study* and *within-cell-type* z-scores exceeded one standard deviation above the mean (z > 1). Cell type–trait categories meeting this criterion were marked with a “+” (Fig. 5).

### MAGMA-based Polygenic Enrichment with scDRS

scDRS is a tool that can assess enrichment of GWAS putative disease genes in an individual cell based on its gene expression profile. ^49^ It relies on MAGMA (Multi-marker Analysis of GenoMic Annotation) for scoring genes from GWAS summary statistics. ^91^ and extracts the top 1000 genes. The polygenic enrichment score per cell represents the aggregated expression of putative disease genes. Leveraging their 74 pre-curated disease/trait MAGMA gene lists, we extracted terms related to the brain and age. We supplemented the selected list with an external MAGMA gene list for PD ^92^, resulting in a total of 25 traits to be tested. For MAGMA, we first annotated genes with SNPs using g1000_eur.bim file for SNP location and NCBI37.3/NCBI37.3.gene.loc file for gene location, both files based on Build 37 (hg19) and downloaded from Complex Trait Genetics Lab https://cncr.nl/research/magma/. Gene window size was set to 10 kb to match scDRS gene sets. This was followed by scoring each gene based on SNPs enrichment of a particular trait and the top 1000 genes were extracted. A gene set file compatible with scDRS was constructed using scdrs munge-gs function. To estimate disease enrichment per cell, we included the number of counts and features per cell as covariates using scdrs compute-score function with default parameters.

### Cosine Similarity

Nuclei were down-sampled to 500 per region. Using PCA space, cosine similarity was calculated between all nuclei-pairs using cosine function from lsa package. This was followed by generating a hierarchically clustered heatmap for visualization of the region-grouped cosine similarities using the Heatmap function from the ComplexHeatmap package.

### Spatial Transcriptomics with 10X Xenium In-Situ Expression Assay

For high-plex spatial transcriptomics, we analyzed tissue sections from rhesus macaque (Macaca mulatta, N=2) and cynomolgus macaque (Macaca fascicularis, N=3). For each animal, we selected 5-7 coronal sections spanning the R-C axis of the striatum.

#### Tissue Preparation

Animals were perfused with 1X PBS and 4% PFA then fixed in sucrose gradients (10% and 20% in4%PFA, and 30% in PBS). After cryoprotection, brains were blocked in the coronal plane, embedded in OCT, frozen,and kept at −80°C for at least 48 h prior to sectioning. Cryosectioning was performed at 20μm and mounted on Xenium Slides (10X Genomics, Cat# 1000660). Slides were stored at −80°C for at least 16 h before proceeding with Xenium Analysis.

#### 10X Genomics Xenium *In Situ* Gene Expression Protocol with Cell Segmentation

(CG000580 Rev E,^93^ CG000749 Rev B^94^): The protocol was performed using Xenium Sample Prep Reagents (10X Genomics, Cat# 1000633), Xenium Cell Segmentation Add-on Kit (10X Genomics, Cat# 1000662), Xenium Decoding Reagents (10X Genomics, Cat# 1000461) and Xenium Decoding Consumables (10X Genomics, Cat# 1000467). Cell segmentation included nuclear expansion (DAPI), boundary segmentation (ATP1A1/CD45/E-Cadherin), and interior segmentation markers—RNA (18S) and protein (alphaSMA/Vimentin). The Xenium gene panel consisted of pre-designed human brain panel (266 genes) supplemented with a custom designed human panel (100 genes). The custom set included 25 marker genes for MSNs cell types and other genes of interest. Imaging, decoding fluorescent probe hybridization and transcript detection were performed on Xenium Analyzer (10X Genomics, PN 1000529) and Xenium Analysis Computer (10X Genomics, Model 1000534). Visualization of key marker genes was carried out using the 10x Xenium Explorer Platform. Downstream analysis including clustering and cell type identification was performed using Seurat v5.

#### Data Processing and Multi-Step Cell Type Identification

##### First Round: Regional Anatomical Identification

The striatum was anatomically identified guided by the spatial localization of *DRD1* and *DRD2* transcript. This dissection was validated with spatial niche assay (BuildNicheAssay function), which uses clusters or initial class annotations to identify cells with similar microenvironments based on the composition of spatially adjacent cell types. K-means clustering grouped cells into spatial niches, enabling precise anatomical dissection of brain regions within the tissue section.

##### Second Round: Striatum Cropping and Filtering

The identified striatum region was cropped based on coordinates, and cells with at least one detected transcript were filtered. Normalization and variance stabilization were performed using the SCTransform() function.

##### Multi-Step Label Transfer

Cell type identification was conducted in a multi-step manner: major cell classes were identified first, followed by subsetting of medium spiny neurons (MSNs) for subtype annotation. Xenium provides subcellular resolution for transcript detection, which facilitates label transfer from snRNA-seq reference using FindTransferAnchors and TransferData functions. However, we noted discrepancies between the observed gene expression and predicted labels for most subtypes. This could be attributed to the cell type heterogeneity in sections across the R-C. This heterogeneity was not fully captured upon relying on labeling from an integrated snRNA-seq reference, where technical differences, as well as real biological-regional differences are mitigated. To address this, we applied Robust Cell Type Decomposition (RCTD), a computational approach for deconvolving spatial pixels using a striatum snRNA-seq reference by estimating the proportions of different cell types contributing to a given spot (create.RCTD() and run.RCTD()). Reference and SpatialRNA objects were created by extracting counts, cluster, and spot information from query and reference objects. Doublet mode was applied to identify pixels as singlets, doublets certain, doublets uncertain and rejected. Pixels labeled as rejected were excluded from analysis, while remaining pixels were annotated according to the cell type with weight >0.5. Major class annotations were then used to build spatial niches with k-means clustering (k=30), which further refined the dissection of the striatal region by excluding off-target regions. The spatial query was filtered to the niche with the highest proportion of MSNs (striatum), followed by dimensionality reduction (PCA, UMAP) and clustering for visualization. For MSN subclass annotation, MSNs were isolated and subtyped using the same label transfer and decomposition workflow. All analyses were conducted using Seurat v5.

### Spatial Regression Discontinuity Design

To evaluate whether novel MSNs subtypes are distinct cell types or instead represent transient cell states within another cell type, we tested whether the expression of certain genes changes abruptly across a spatial boundary in the tissue. In particular, we assessed whether transitions in a marker gene expression across a spatial axis is discrete or continuous within a pair of related cell types. We applied a regression discontinuity design ^95^, modeling gene expression as a function of spatial position and testing whether introducing a threshold along the spatial axis explains additional variance. The null hypothesis was that the gene expression is a linear function of the spatial location, and adding a threshold does not add additional information. First we defined the spatial axis as x- or y-coordinates independently to test for gene expression discontinuity along either direction. Alternatively, we derived a non-linear 1D spatial embedding using isometric mapping (sklearn.manifold.Isomap) to infer a non-linear 1D spatial embedding that represents the anatomical gradient and the local arrangement of different cell populations. Second, we used Otsu thresholding to identify a spatial boundary that partitions the spatial axis into two regions and then centered the spatial axis coordinates along this threshold (boundary = 0), where cells that lie to the right of the threshold boundary are labeled as positive and on the left are labeled as negative. Third, we fit a gaussian generalized linear model (GLM) of the form: *smf.glm(formula=“gene ∼ posmark + centered_axis”, family=sm.families.Gaussian())*, to test whether the gene expression differs significantly across the spatial threshold, while controlling for continuous spatial position. To confirm biological reproducibility, we repeated this test on two independent sections from different rhesus macaques. Specifically, we examined *TAC3* expression shift in D1-VS-RGS6 and D1-VS to evaluate whether D1-VS-RGS6 – previously identified as *TAC3* archetype within the D1-shell/OT subtype ^26^ – represents a distinct subtype or a subcluster within D1-VS. This analysis revealed a significant spatial discontinuity in *TAC3* expression supporting the identification of D1-VS-RGS6 as distinct MSN subtype.

## CODE AND DATA AVAILABILITY

Data and code will be available at GEO and ASAP CRN Cloud at the time of publication. All R and Python scripts used for data processing, analysis and generation of main and supplementary figures are available at https://github.com/pfenninglab/ and https://github.com/rewardlab/. Any additional data, custom code or resources not included in the repository can be requested from the corresponding author.

## ACKNOWLEDGMENTS

The authors thank Jaquelyn Breter for animal care and enrichment and Mitsutoshi Hanada, Natasha Nosenchuck, Nyona Towler, Aydin Alikaya, Nili Shah for assistance with brain tissue embedding, cutting, and assistance with spatial transcriptomic tissue processing. This work was supported by NIH grants UG3MH120094 and UF1MH130881 (A.R.P. and W.R.S.), R01NS100908 and P51OD011132 (A.G.), and by Aligning Science Across Parkinson’s (ASAP-020519, ASAP-025193) (W.R.S.) through the Michael J. Fox Foundation for Parkinson’s Research (MJFF). For the purposes of open access, the authors have applied a CC-BY public copyright license to all Author Accepted Manuscripts arising from this submission.

## Notes

### Competing Interest Statement

The authors have declared no competing interest.

### Summary of Updates

This version includes the results section associated with Figure 4.

## REFERENCES

1. Samejima, K., Ueda, Y., Doya, K., and Kimura, M. (2005). Representation of action-specific reward values in the striatum. Science 310, 1337–1340.

2. Alexander, G.E., and DeLong, M.R. (1985). Microstimulation of the primate neostriatum. I. Physiological properties of striatal microexcitable zones. J. Neurophysiol. 53, 1401–1416.

3. Alexander, G.E., and DeLong, M.R. (1985). Microstimulation of the primate neostriatum. II. Somatotopic organization of striatal microexcitable zones and their relation to neuronal response properties. J. Neurophysiol. 53, 1417–1430.

4. Hikosaka, O., and Sakamoto, M. (1986). Neural activities in the monkey basal ganglia related to attention, memory and anticipation. Brain Dev. 8, 454–461.

5. Ding, L., and Gold, J.I. (2013). The basal ganglia’s contributions to perceptual decision making. Neuron 79, 640–649.

6. Hikosaka, O., Takikawa, Y., and Kawagoe, R. (2000). Role of the basal ganglia in the control of purposive saccadic eye movements. Preprint, https://doi.org/10.1152/physrev.2000.80.3.953 10.1152/physrev.2000.80.3.953.

7. DeLong, M.R. (1990). Primate models of movement disorders of basal ganglia origin. 13, 281–285.

8. Stein, D.J. (2002). Obsessive-compulsive disorder. Lancet 360, 397–405.

9. Volkow, N.D., and Morales, M. (2015). The brain on drugs: From reward to addiction. Cell 162, 712–725.

10. Alexander, G.E., DeLong, M.R., and Strick, P.L. (1986). Parallel organization of functionally segregated circuits linking basal ganglia and cortex. Annu. Rev. Neurosci. 9, 357–381.

11. Kemp, J.M., and Powell, T.P. (1970). The cortico-striate projection in the monkey. Brain 93, 525–546.

12. Gerfen, C.R., Engber, T.M., Mahan, L.C., Susel, Z., Chase, T.N., Monsma, F.J., Jr, and Sibley, D.R. (1990). D1 and D2 dopamine receptor-regulated gene expression of striatonigral and striatopallidal neurons. Science 250, 1429–1432.

13. Gong, S., Doughty, M., Harbaugh, C.R., Cummins, A., Hatten, M.E., Heintz, N., and Gerfen, C.R. (2007). Targeting Cre recombinase to specific neuron populations with bacterial artificial chromosome constructs. J. Neurosci. 27, 9817–9823.

14. Gerfen, C.R., Paletzki, R., and Heintz, N. (2013). GENSAT BAC cre-recombinase driver lines to study the functional organization of cerebral cortical and basal ganglia circuits. Neuron 80, 1368–1383.

15. Lazaridis, I., Crittenden, J.R., Ahn, G., Hirokane, K., Wickersham, I.R., Yoshida, T., Mahar, A., Skara, V., Loftus, J.H., Parvataneni, K., et al. (2024). Striosomes control dopamine via dual pathways paralleling canonical basal ganglia circuits. Curr. Biol. 34, 5263–5283.e8.

16. Yao, Z., van Velthoven, C.T.J., Kunst, M., Zhang, M., McMillen, D., Lee, C., Jung, W., Goldy, J., Abdelhak, A., Aitken, M., et al. (2023). A high-resolution transcriptomic and spatial atlas of cell types in the whole mouse brain. Nature 624, 317–332.

17. Zhou, J., Zhang, Z., Wu, M., Liu, H., Pang, Y., Bartlett, A., Peng, Z., Ding, W., Rivkin, A., Lagos, W.N., et al. (2023). Brain-wide correspondence of neuronal epigenomics and distant projections. Nature 624, 355–365.

18. Xiong, X., James, B.T., Boix, C.A., Park, Y.P., Galani, K., Victor, M.B., Sun, N., Hou, L., Ho, L.L., Mantero, J., et al. (2023). Epigenomic dissection of Alzheimer’s disease pinpoints causal variants and reveals epigenome erosion. Cell. 10.1016/j.cell.2023.08.040.

19. Liu, Z., Zhang, S., James, B.T., Galani, K., Mangan, R.J., Fass, S.B., Liang, C., Wagle, M.M., Boix, C.A., Tanigawa, Y., et al. (2025). Single-cell multiregion epigenomic rewiring in Alzheimer’s disease progression and cognitive resilience. Cell 188, 4980–5002.e29.

20. Mathys, H., Peng, Z., Boix, C.A., Victor, M.B., Leary, N., Babu, S., Abdelhady, G., Jiang, X., Ng, A.P., Ghafari, K., et al. (2023). Single-cell atlas reveals correlates of high cognitive function, dementia, and resilience to Alzheimer’s disease pathology. Cell 186, 4365–4385.e27.

21. Zemke, N.R., Armand, E.J., Wang, W., Lee, S., Zhou, J., Li, Y.E., Liu, H., Tian, W., Nery, J.R., Castanon, R.G., et al. (2023). Conserved and divergent gene regulatory programs of the mammalian neocortex. Nature 624, 390–402.

22. Izpisua Belmonte, J.C., Callaway, E.M., Caddick, S.J., Churchland, P., Feng, G., Homanics, G.E., Lee, K.F., Leopold, D.A., Miller, C.T., Mitchell, J.F., et al. (2015). Brains, genes, and primates. Neuron 86, 617–631.

23. Wichmann, T., and DeLong, M.R. (2006). Deep brain stimulation for neurologic and neuropsychiatric disorders. Neuron 52, 197–204.

24. Bronstein, J.M., Tagliati, M., Alterman, R.L., Lozano, A.M., Volkmann, J., Stefani, A., Horak, F.B., Okun, M.S., Foote, K.D., Krack, P., et al. (2011). Deep brain stimulation for Parkinson disease: an expert consensus and review of key issues. 68, 165–165.

25. Bergman, H., Wichmann, T., and DeLong, M.R. (1990). Reversal of experimental parkinsonism by lesions of the subthalamic nucleus. Science 249, 1436–1438.

26. He, J., Kleyman, M., Chen, J., Alikaya, A., Rothenhoefer, K.M., Ozturk, B.E., Wirthlin, M., Bostan, A.C., Fish, K., Byrne, L.C., et al. (2021). Transcriptional and anatomical diversity of medium spiny neurons in the primate striatum. Curr. Biol. 10.1016/j.cub.2021.10.015.

27. Krienen, F.M., Goldman, M., Zhang, Q., C H Del Rosario, R., Florio, M., Machold, R., Saunders, A., Levandowski, K., Zaniewski, H., Schuman, B., et al. (2020). Innovations present in the primate interneuron repertoire. Nature 586, 262–269.

28. Märtin, A., Calvigioni, D., Tzortzi, O., Fuzik, J., Wärnberg, E., and Meletis, K. (2019). A Spatiomolecular Map of the Striatum. Cell Rep. 29, 4320–4333 e5.

29. Gerfen, C.R. (1989). The neostriatal mosaic: striatal patch-matrix organization is related to cortical lamination. Science 246, 385–388.

30. Gerfen, C.R., Herkenham, M., and Thibault, J. (1987). The neostriatal mosaic: II. Patch-and matrix-directed mesostriatal dopaminergic and non-dopaminergic systems. J. Neurosci. 7, 3915–3934.

31. Graybiel, A.M., and Ragsdale, C.W., Jr (1978). Histochemically distinct compartments in the striatum of human, monkeys, and cat demonstrated by acetylthiocholinesterase staining. Proc Natl Acad Sci U S A 75, 5723–5726.

32. Fudge, J.L., and Haber, S.N. (2002). Defining the caudal ventral striatum in primates: cellular and histochemical features. J. Neurosci. 22, 10078–10082.

33. Cable, D.M., Murray, E., Zou, L.S., Goeva, A., Macosko, E.Z., Chen, F., and Irizarry, R.A. (2022). Robust decomposition of cell type mixtures in spatial transcriptomics. Nat. Biotechnol. 40, 517–526.

34. Del Rey, N.L.-G., Trigo-Damas, I., Obeso, J.A., Cavada, C., and Blesa, J. (2022). Neuron types in the primate striatum: Stereological analysis of projection neurons and interneurons in control and parkinsonian monkeys. Neuropathol. Appl. Neurobiol. 48, e12812.

35. Graveland, G.A., and DiFiglia, M. (1985). The frequency and distribution of medium-sized neurons with indented nuclei in the primate and rodent neostriatum. Brain Res. 327, 307–311.

36. Ahlers, K.E., Chakravarti, B., and Fisher, R.A. (2016). RGS6 as a novel therapeutic target in CNS diseases and cancer. Preprint, https://doi.org/10.1208/s12248-016-9899-9 10.1208/s12248-016-9899-9.

37. Srinivasan, C., Phan, B.N., Lawler, A.J., Ramamurthy, E., Kleyman, M., Brown, A.R., Kaplow, I.M., Wirthlin, M.E., and Pfenning, A.R. (2021). Addiction-associated genetic variants implicate brain cell type- and region-specific Cis-regulatory elements in addiction neurobiology. J. Neurosci. 41, 9008–9030.

38. Corces, M.R., Shcherbina, A., Kundu, S., Gloudemans, M.J., Frésard, L., Granja, J.M., Louie, B.H., Eulalio, T., Shams, S., Bagdatli, S.T., et al. (2020). Single-cell epigenomic analyses implicate candidate causal variants at inherited risk loci for Alzheimer’s and Parkinson’s diseases. Nat Genet 52, 1158–1168.

39. Bryois, J., Skene, N.G., Hansen, T.F., Kogelman, L.J.A., Watson, H.J., Liu, Z., Eating Disorders Working Group of the Psychiatric Genomics Consortium, International Headache Genetics Consortium, 23andMe Research Team, Brueggeman, L., et al. (2020). Genetic identification of cell types underlying brain complex traits yields insights into the etiology of Parkinson’s disease. Nat. Genet. 52, 482–493.

40. Skene, N.G., Bryois, J., Bakken, T.E., Breen, G., Crowley, J.J., Gaspar, H.A., Giusti-Rodriguez, P., Hodge, R.D., Miller, J.A., Muñoz-Manchado, A.B., et al. (2018). Genetic identification of brain cell types underlying schizophrenia. Nat. Genet. 50, 825–833.

41. Gayden, J., Puig, S., Srinivasan, C., Phan, B.N., Abdelhady, G., Buck, S.A., Gamble, M.C., Tejeda, H.A., Dong, Y., Pfenning, A.R., et al. (2023). Integrative multi-dimensional characterization of striatal projection neuron heterogeneity in adult brain. bioRxivorg, 2023.05.04.539488.

42. Ferreira, P.G., Muñoz-Aguirre, M., Reverter, F., Sá Godinho, C.P., Sousa, A., Amadoz, A., Sodaei, R., Hidalgo, M.R., Pervouchine, D., Carbonell-Caballero, J., et al. (2018). The effects of death and post-mortem cold ischemia on human tissue transcriptomes. Nat. Commun. 9, 490.

43. Bulik-Sullivan, B.K., and Neale, B.M. (2015). LD Score Regression Distinguishes Confounding from Polygenicity in GWAS. Nat. Genet.

44. Finucane, H.K., Bulik-Sullivan, B., Gusev, A., Trynka, G., Reshef, Y., Loh, P.R., Anttila, V., Xu, H., Zang, C., Farh, K., et al. (2015). Partitioning heritability by functional annotation using genome-wide association summary statistics. Nat. Genet. 10.1038/ng.3404.

45. Akiyama, H., Barger, S., Barnum, S., Bradt, B., Bauer, J., Cole, G.M., Cooper, N.R., Eikelenboom, P., Emmerling, M., Fiebich, B.L., et al. (2000). Inflammation and Alzheimer’s disease. Neurobiol. Aging 21, 383–421.

46. McCutcheon, R.A., Abi-Dargham, A., and Howes, O.D. (2019). Schizophrenia, dopamine and the striatum: From biology to symptoms. Trends Neurosci. 42, 205–220.

47. Savage, J.E., Jansen, P.R., Stringer, S., Watanabe, K., Bryois, J., de Leeuw, C.A., Nagel, M., Awasthi, S., Barr, P.B., Coleman, J.R.I., et al. (2018). Genome-wide association meta-analysis in 269,867 individuals identifies new genetic and functional links to intelligence. Nat. Genet. 50, 912–919.

48. Lee, J.J., Wedow, R., Okbay, A., Kong, E., Maghzian, O., Zacher, M., Nguyen-Viet, T.A., Bowers, P., Sidorenko, J., Karlsson Linnér, R., et al. (2018). Gene discovery and polygenic prediction from a genome-wide association study of educational attainment in 1.1 million individuals. Nat. Genet. 50, 1112–1121.

49. Zhang, M.J., Hou, K., Dey, K.K., Sakaue, S., Jagadeesh, K.A., Weinand, K., Taychameekiatchai, A., Rao, P., Pisco, A.O., Zou, J., et al. (2022). Polygenic enrichment distinguishes disease associations of individual cells in single-cell RNA-seq data. Nat. Genet. 54, 1572–1580.

50. Young, W.S., 3rd, Bonner, T.I., and Brann, M.R. (1986). Mesencephalic dopamine neurons regulate the expression of neuropeptide mRNAs in the rat forebrain. Proc. Natl. Acad. Sci. U. S. A. 83, 9827–9831.

51. Albin, R.L., Young, A.B., and Penney, J.B. (1989). The functional anatomy of basal ganglia disorders. Trends Neurosci. 10.1016/0166-2236(89)90074-X.

52. DeLong, M.R., and Wichmann, T. (2007). Circuits and circuit disorders of the basal ganglia. Arch. Neurol. 64, 20–24.

53. Harris, J.P., Burrell, J.C., Struzyna, L.A., Chen, H.I., Serruya, M.D., Wolf, J.A., Duda, J.E., and Cullen, D.K. (2020). Emerging regenerative medicine and tissue engineering strategies for Parkinson’s disease. Preprint, https://doi.org/10.1038/s41531-019-0105-5, 10.1038/s41531-019-0105-5.

54. Chen, Y., Hong, Z., Wang, J., Liu, K., Liu, J., Lin, J., Feng, S., Zhang, T., Shan, L., Liu, T., et al. (2023). Circuit-specific gene therapy reverses core symptoms in a primate Parkinson’s disease model. Cell. 10.1016/j.cell.2023.10.004.

55. Flajolet, M., Wang, Z., Futter, M., Shen, W., Nuangchamnong, N., Bendor, J., Wallach, I., Nairn, A.C., Surmeier, D.J., and Greengard, P. (2008). FGF acts as a co-transmitter through adenosine A2A receptor to regulate synaptic plasticity. Nat. Neurosci. 10.1038/nn.2216.

56. Luo, Z., Ahlers-Dannen, K.E., Spicer, M.M., Yang, J., Alberico, S., Stevens, H.E., Narayanan, N.S., and Fisher, R.A. (2019). Age-dependent nigral dopaminergic neurodegeneration and α-synuclein accumulation in RGS6-deficient mice. JCI Insight. 10.1172/jci.insight.126769.

57. Ahlers-Dannen, K.E., Spicer, M.M., and Fisher, R.A. (2020). RGS proteins as critical regulators of motor function and their implications in Parkinson’s disease. Mol. Pharmacol. 10.1124/MOL.119.118836.

58. Bifsha, P., Yang, J., Fisher, R.A., and Drouin, J. (2014). Rgs6 is Required for Adult Maintenance of Dopaminergic Neurons in the Ventral Substantia Nigra. PLoS Genet. 10.1371/journal.pgen.1004863.

59. Geurts, M., Maloteaux, J.M., and Hermans, E. (2003). Altered expression of regulators of G-protein signaling (RGS) mRNAs in the striatum of rats undergoing dopamine depletion. Biochem. Pharmacol. 10.1016/S0006-2952(03)00447-7.

60. Kovoor, A., Seyffarth, P., Ebert, J., Barghshoon, S., Chen, C.K., Schwarz, S., Axelrod, J.D., Cheyette, B.N.R., Simon, M.I., Lester, H.A., et al. (2005). D2 dopamine receptors colocalize regulator of G-protein signaling 9-2 (RGS9-2) via the RGS9 DEP domain, and RGS9 knock-out mice develop dyskinesias associated with dopamine pathways. J. Neurosci. 10.1523/JNEUROSCI.2840-04.2005.

61. Miyamoto, Y., Katayama, S., Shigematsu, N., Nishi, A., and Fukuda, T. (2018). Striosome-based map of the mouse striatum that is conformable to both cortical afferent topography and uneven distributions of dopamine D1 and D2 receptor-expressing cells. Brain Struct. Funct. 10.1007/s00429-018-1749-3.

62. Van Hoesen, G.W., Yeterian, E.H., and Lavizzo-Mourey, R. (1981). Widespread corticostriate projections from temporal cortex of the rhesus monkey. J. Comp. Neurol. 199, 205–219.

63. Yeterian, E.H., and Van Hoesen, G.W. (1978). Cortico-striate projections in the rhesus monkey: the organization of certain cortico-caudate connections. Brain Res. 139, 43–63.

64. Kunimatsu, J., Maeda, K., and Hikosaka, O. (2019). The caudal part of putamen represents the historical object value information. J. Neurosci. 39, 1709–1719.

65. Amita, H., Kim, H.F., Smith, M.K., Gopal, A., and Hikosaka, O. (2019). Neuronal connections of direct and indirect pathways for stable value memory in caudal basal ganglia. Eur. J. Neurosci. 49, 712–725.

66. He, J., Phan, B.N., Kerkhoff, W.G., Alikaya, A., Hong, T., Brull, O.R., Fredericks, J.M., Sedorovitz, M., Srinivasan, C., Leone, M.J., et al. (2025). Machine learning identification of enhancers in the rhesus macaque genome. Neuron 113, 1548–1561.e8.

67. Maurano, M.T., Humbert, R., Rynes, E., Thurman, R.E., Haugen, E., Wang, H., Reynolds, A.P., Sandstrom, R., Qu, H., Brody, J., et al. (2012). Systematic localization of common disease-associated variation in regulatory DNA. Science 337, 1190–1195.

68. McQuade, A., and Blurton-Jones, M. (2019). Microglia in Alzheimer’s disease: Exploring how genetics and phenotype influence risk. J. Mol. Biol. 431, 1805–1817.

69. Kosoy, R., Fullard, J.F., Zeng, B., Bendl, J., Dong, P., Rahman, S., Kleopoulos, S.P., Shao, Z., Girdhar, K., Humphrey, J., et al. (2022). Genetics of the human microglia regulome refines Alzheimer’s disease risk loci. Nat. Genet. 54, 1145–1154.

70. Logue, M.W., Schu, M., Vardarajan, B.N., Farrell, J., Lunetta, K.L., Jun, G., Baldwin, C.T., Deangelis, M.M., and Farrer, L.A. (2014). Search for age-related macular degeneration risk variants in Alzheimer disease genes and pathways. Neurobiol. Aging 35, 1510.e7–e18.

71. Trubetskoy, V., Pardiñas, A.F., Qi, T., Panagiotaropoulou, G., Awasthi, S., Bigdeli, T.B., Bryois, J., Chen, C.-Y., Dennison, C.A., Hall, L.S., et al. (2022). Mapping genomic loci implicates genes and synaptic biology in schizophrenia. Nature 604, 502–508.

72. Pardiñas, A.F., Holmans, P., Pocklington, A.J., Escott-Price, V., Ripke, S., Carrera, N., Legge, S.E., Bishop, S., Cameron, D., Hamshere, M.L., et al. (2018). Common schizophrenia alleles are enriched in mutation-intolerant genes and in regions under strong background selection. Nat. Genet. 50, 381–389.

73. Warren, W.C., Harris, R.A., Haukness, M., Fiddes, I.T., Murali, S.C., Fernandes, J., Dishuck, P.C., Storer, J.M., Raveendran, M., Hillier, L.D.W., et al. (2020). Sequence diversity analyses of an improved rhesus macaque genome enhance its biomedical utility. Science. 10.1126/science.abc6617.

74. Young, M.D., and Behjati, S. (2020). SoupX removes ambient RNA contamination from droplet-based single-cell RNA sequencing data. Gigascience. 10.1093/gigascience/giaa151.

75. McGinnis, C.S., Murrow, L.M., and Gartner, Z.J. (2019). DoubletFinder: Doublet Detection in Single-Cell RNA Sequencing Data Using Artificial Nearest Neighbors. Cell Syst. 10.1016/j.cels.2019.03.003.

76. Hao, Y., Stuart, T., Kowalski, M.H., Choudhary, S., Hoffman, P., Hartman, A., Srivastava, A., Molla, G., Madad, S., Fernandez-Granda, C., et al. (2024). Dictionary learning for integrative, multimodal and scalable single-cell analysis. Nat. Biotechnol. 42, 293–304.

77. Law, C.W., Chen, Y., Shi, W., and Smyth, G.K. (2014). voom: Precision weights unlock linear model analysis tools for RNA-seq read counts. Genome Biol. 15, R29.

78. Smyth, G.K. (2005). limma: Linear Models for Microarray Data. In Bioinformatics and Computational Biology Solutions Using R and Bioconductor (Springer-Verlag), pp. 397–420.

79. Phan, B.D.N., Ray, M.H., Xue, X., Fu, C., Fenster, R.J., Kohut, S.J., Bergman, J., Haber, S.N., McCullough, K.M., Fish, M.K., et al. (2024). Single nuclei transcriptomics in human and non-human primate striatum in opioid use disorder. Nat. Commun. 10.1038/s41467-024-45165-7.

80. Boca, S.M., and Leek, J.T. (2018). A direct approach to estimating false discovery rates conditional on covariates. PeerJ 6, e6035.

81. Zhang, H., Song, L., Wang, X., Cheng, H., Wang, C., Meyer, C.A., Liu, T., Tang, M., Aluru, S., Yue, F., et al. (2021). Fast alignment and preprocessing of chromatin profiles with Chromap. Nat. Commun. 12, 6566.

82. Shumate, A., and Salzberg, S.L. (2021). Liftoff: accurate mapping of gene annotations. Bioinformatics 37, 1639–1643.

83. Granja, J.M., Corces, M.R., Pierce, S.E., Bagdatli, S.T., Choudhry, H., Chang, H.Y., and Greenleaf, W.J. (2021). ArchR is a scalable software package for integrative single-cell chromatin accessibility analysis. Nat. Genet. 10.1038/s41588-021-00790-6.

84. Bon, J.J., Bretherton, A., Buchhorn, K., Cramb, S., Drovandi, C., Hassan, C., Jenner, A.L., Mayfield, H.J., McGree, J.M., Mengersen, K., et al. (2023). Being Bayesian in the 2020s: opportunities and challenges in the practice of modern applied Bayesian statistics. Philos. Trans. A Math. Phys. Eng. Sci. 381, 20220156.

85. Korsunsky, I., Millard, N., Fan, J., Slowikowski, K., Zhang, F., Wei, K., Baglaenko, Y., Brenner, M., Loh, P.-R., and Raychaudhuri, S. (2019). Fast, sensitive and accurate integration of single-cell data with Harmony. Nat. Methods 16, 1289–1296.

86. Zhang, Y., Liu, T., Meyer, C.A., Eeckhoute, J., Johnson, D.S., Bernstein, B.E., Nusbaum, C., Myers, R.M., Brown, M., Li, W., et al. (2008). Model-based analysis of ChIP-Seq (MACS). Genome Biol. 9, R137.

87. Finucane, H.K., Reshef, Y.A., Anttila, V., Slowikowski, K., Gusev, A., Byrnes, A., Gazal, S., Loh, P.-R., Lareau, C., Shoresh, N., et al. (2018). Heritability enrichment of specifically expressed genes identifies disease-relevant tissues and cell types. Nat. Genet. 50, 621–629.

88. Tashman, K.C., Cui, R., O’Connor, L.J., Neale, B.M., and Finucane, H.K. (2021). Significance testing for small annotations in stratified LD-Score regression. Preprint.

89. Armstrong, J., Hickey, G., Diekhans, M., Fiddes, I.T., Novak, A.M., Deran, A., Fang, Q., Xie, D., Feng, S., Stiller, J., et al. (2020). Progressive Cactus is a multiple-genome aligner for the thousand-genome era. Nature. 10.1038/s41586-020-2871-y.

90. Zhang, X., Kaplow, I.M., Wirthlin, M., Park, T.Y., and Pfenning, A.R. (2020). HALPER facilitates the identification of regulatory element orthologs across species. Comput. Appl. Biosci. 10.1093/bioinformatics/btaa493.

91. de Leeuw, C.A., Mooij, J.M., Heskes, T., and Posthuma, D. (2015). MAGMA: generalized gene-set analysis of GWAS data. PLoS Comput. Biol. 11, e1004219.

92. Nalls, M.A., Blauwendraat, C., Vallerga, C.L., Heilbron, K., Bandres-Ciga, S., Chang, D., Tan, M., Kia, D.A., Noyce, A.J., Xue, A., et al. (2019). Identification of novel risk loci, causal insights, and heritable risk for Parkinson’s disease: a meta-analysis of genome-wide association studies. Lancet Neurol. 18, 1091–1102.

93. 10x Genomics (2024). Xenium In Situ for FFPE – Deparaffinization & Decrosslinking Demonstrated Protocol.

94. 10x Genomics (2024). Xenium In Situ Gene Expression with Cell Segmentation Staining User Guide.

95. Bor, J., Moscoe, E., Mutevedzi, P., Newell, M.-L., and Bärnighausen, T. (2014). Regression Discontinuity Designs in Epidemiology. Epidemiology. 10.1097/ede.0000000000000138.

